# Healthy ageing reduces the precision of episodic memory retrieval

**DOI:** 10.1101/468579

**Authors:** Saana M. Korkki, Franziska R. Richter, Priyanga Jeyarathnarajah, Jon S. Simons

## Abstract

Episodic memory declines with older age, but it is unresolved whether this decline reflects reduced probability of successfully retrieving information from memory, or decreased precision of the retrieved information. Here, we used continuous measures of episodic memory retrieval in combination with computational modelling of participants’ retrieval errors to distinguish between these two potential accounts of age-related memory deficits. In three experiments, young and older participants encoded stimuli displays consisting of everyday objects varying along different perceptual features (e.g., location, colour and orientation) in a circular space. At test, participants recreated the features of studied objects using a continuous response dial. Across all three experiments, we observed age-related declines in the precision of episodic memory retrieval, whereas age differences in retrieval success were limited to the most challenging task condition. Reductions in mnemonic precision were evident for retrieval of both item-based and contextual information, and persisted after controlling for age-related decreases in the fidelity of perception and working memory. The findings highlight impoverished precision of memory representations as one factor contributing to age-related episodic memory loss, and suggest that the cognitive and neural changes associated with older age can differentially affect distinct aspects of episodic retrieval.

---

Episodic memory enables us to recollect details of events from our personal pasts, such as recalling our last birthday party, or where we parked our car on our last visit to the supermarket. Intact memories of our past experiences are vital for developing and maintaining our sense of self (Conway, 2005; Tulving, 2002), and guide the actions and decisions we take in our everyday lives (Schacter, Addis, & Buckner, 2007; Wimmer & Shohamy, 2012), enabling flexible behaviour in changing environments. Episodic memory function exhibits marked declines as we grow older (Grady, 2012; Hedden & Gabrieli, 2004), however, with longitudinal studies typically displaying decreases from the age of 60 onward (Nyberg, Lövdén, Riklund, Lindenberger, & Bäckman, 2012; Nyberg & Pudas, 2019). The particular vulnerability of episodic memory to age-related decline in comparison to other cognitive domains, including other types of long-term memory, has been highlighted in previous studies (Nyberg et al., 2003; Rönnlund, Nyberg, Bäckman, & Nilsson, 2005), but the specific neurocognitive mechanisms underlying this impairment are yet to be fully characterised. In particular, it is unclear whether age-related memory reductions reflect a decreased probability of successfully retrieving information from memory, or more qualitative changes in the fidelity with which memory traces can be encoded into and retrieved from memory.

In typical laboratory tests of episodic memory, participants’ performance is measured using categorical response options, for example by asking a participant to judge whether a test stimulus has been previously encountered (“old”) or not (“new”). These types of measures, however, often afford only binary distinctions between successful and unsuccessful memory retrieval, unable to fully capture the multifaceted nature of episodic recollection. Increasing evidence suggests that instead of an “all-or-none” process, varying only in the dichotomy between successful and unsuccessful retrieval, episodic recollection likely operates in a “some-or-none” manner, where the quality, or precision, of the successfully retrieved information can vary on a graded scale (Harlow & Donaldson, 2013; Onyper, Zhang, & Howard, 2010; Yonelinas & Parks, 2007). To investigate these more fine-grained variations in episodic memory, recent studies have begun to utilize continuous measures of retrieval performance, where participants are asked to reconstruct aspects of the studied stimuli using a continuous, analogue scale. In younger adults, studies employing these types of tasks have demonstrated retrieval success and precision to be separable components of long-term memory (LTM) (Harlow & Donaldson, 2013; Harlow & Yonelinas, 2016; Richter, Cooper, Bays, & Simons, 2016), which can be selectively affected by experimental manipulations (e.g., Sutterer & Awh, 2016; Xie & Zhang, 2017), brain stimulation (Nilakantan, Bridge, Gagnon, VanHaerents, & Voss, 2017), and developmental condition (Cooper et al., 2017). Furthermore, a recent study by Richter, Cooper and colleagues (2016) provided evidence for a dissociation between these two mnemonic constructs at the neural level, demonstrating that the success and precision of episodic recollection rely on distinct brain regions of the core memory network, with the probability of successful memory retrieval scaling with hippocampal activity and the precision of memory retrieval with activity in the angular gyrus. Given the dissociable neurocognitive profiles of these two subcomponents of episodic memory retrieval in younger adults, it is also possible that they are differentially sensitive to age-related cognitive decline.

Traditionally, investigations of age-related changes in episodic memory have focused on examining the effects of older age on measures of retrieval success (e.g., Cansino et al., 2018; Craik & McDowd, 1987; Koutstaal, 2003; Mark & Rugg, 1998; Naveh-Benjamin, 2000). However, several strands of evidence imply that memory function in older age might to some extent be constrained by reductions in the quality and specificity of information retained in memory. For example, age-related increases in false memory have been interpreted as resulting from increased reliance on gist-like representations of previous events with diminished encoding and retrieval of specific stimuli details (Dennis, Kim, & Cabeza, 2007, 2008; Kensinger & Schacter, 1999; Koutstaal & Schacter, 1997). Furthermore, previous research has demonstrated greater age differences in episodic recollection when participants are required to retrieve more detailed information about the study event (Luo & Craik, 2009), and that older adults tend to recall less specific details of events from their personal pasts in comparison to younger adults (Addis, Wong, & Schacter, 2008; Levine, Svoboda, Hay, Winocur, & Moscovitch, 2002). Despite often preserved ability to recognise studied items as previously encountered, and to identify dissimilar novel items as new, older adults are also typically impaired in mnemonic discrimination of studied items from perceptually similar lures (Stark, Yassa, Lacy, & Stark, 2013; Toner, Pirogovsky, Kirwan, & Gilbert, 2009; Yassa et al., 2011), implying a reduced level of detail of the retained memory representations in older age. At the neural level, functional brain imaging has further indicated age-related decreases in the fidelity of neural representations corresponding to different stimuli or task contexts during both encoding and retrieval of episodic memory (Abdulrahman, Fletcher, Bullmore, & Morcom, 2017; St-Laurent, Abdi, Bondad, & Buchsbaum, 2014; Trelle, Henson, & Simons, 2018; Zheng et al., 2018), potentially constraining the precision with which memory representations can be formed as well as recovered in older age.

Despite signs of reduced memory quality in ageing, the majority of previous investigations have tended to rely on categorical measures of memory performance, which are unable to discern whether age-related performance reductions are due to changes in the success or precision of memory retrieval. For instance, a failure to correctly retrieve a specific study detail in a categorical memory task could reflect either a failure to access the information in question, or decreased fidelity of the retrieved information, leading to selection of an incorrect retrieval response. In working memory (WM) research, continuous report tasks, providing a more direct measure of memory fidelity, have been fruitful in elucidating the specific components of short-term memory degradation in older age, revealing age-related decreases in mnemonic precision and increases in binding errors, whereas no age differences in the success of memory retrieval were detected (Peich, Husain, & Bays, 2013). This approach has recently been extended to investigate age-related changes in object-spatial location binding in long-term memory, suggesting that the precision of LTM retrieval might similarly be sensitive to age-related decline (Nilakantan, Bridge, VanHaerents, & Voss, 2018).

The aim of the current study was to employ a continuous report paradigm, adapted from recent work in younger adults (Richter, Cooper et al., 2016), to better characterise the nature of age-related changes in episodic memory. Specifically, we aimed to distinguish whether age-related memory decreases reflect reduced probability of successfully retrieving information from memory, and/or decreased precision of the retrieved memory representations. In a series of three experiments, healthy young and older participants encoded visual stimuli displays consisting of everyday objects varying along different perceptual features (e.g., location, colour and orientation) in a circular space. At test, participants were asked to recreate the features of studied objects using a continuous response dial, allowing for detailed assessment of retrieval performance. Fitting a computational model (Bays, Catalao, & Husain, 2009; Zhang & Luck, 2008) to participants’ retrieval error data allowed us to estimate both the probability of successful retrieval and the precision of the retrieved information from the same data, distinguishing between these two alternative sources of memory errors in older age.

Previous research has implicated the retrieval of contextual information, such as spatial location, as particularly sensitive to age-related degradation (Chalfonte & Johnson, 1996; Kukolja, Thiel, Wilms, Mirzazade, & Fink, 2009; Nilakantan et al., 2018; Old & Naveh-Benjamin, 2008; Rajah, Languay, & Valiquette, 2010; Spencer & Raz, 1995), therefore in the first experiment we used a location memory task to examine the effects of ageing on the success and precision of episodic memory retrieval. In contrast to memory for the context in which studied items were encountered, memory for the items themselves has typically been considered less vulnerable to age-related decline (Old & Naveh-Benjamin, 2008; Spencer & Raz, 1995). In the second experiment, we thus examined whether similar deficits in item-based as well as contextual memory could be detected using the continuous report task. In the third experiment, we investigated how specific any age-related changes in the success and precision of episodic memory retrieval were to long-term memory processes, or whether they could be to some extent explained by potential deficits at the level of perception (Monge & Madden, 2016) or working memory (Peich et al., 2013; Pertzov, Heider, Liang, & Husain, 2015).

## General methods

In each experiment, participants encoded object stimuli displays and later recreated the features (such as location, colour, or orientation) of studied objects as precisely as they could using a 360-degree response dial. Both studied feature values and participant responses mapped onto a circular space, enabling us to distinguish between the probability of successful retrieval (i.e., probability of retrieving some information about the correct target feature value) and the precision of retrieved information (i.e., variability in successful target retrieval) with a computational modelling approach derived from working memory research (Bays et al., 2009; Zhang & Luck, 2008), but more recently also applied to long-term memory studies (e.g., Brady, Konkle, Gill, Oliva, & Alvarez, 2013; Richter, Cooper et al., 2016). At the beginning of each experiment, participants completed a demographic questionnaire and the Shipley Institute of Living Vocabulary Scale (SILVS) (Zachary & Shipley, 1986) measure of crystallized intelligence. Older adults additionally completed the Montreal Cognitive Assessment (MoCA) (Nasreddine et al., 2005), a screening tool for mild cognitive impairment. Before each of the continuous report tasks, participants completed practice trials of the task.

### Participants

Participants for all experiments were native English-speakers who reported normal or corrected-to-normal vision, no colour blindness, and no current or historical diagnosis of any psychiatric or neurological condition, or learning difficulty. Older participants scored in the healthy range (26 or above) on the MoCA (Nasreddine et al., 2005). Participants gave written and informed consent in a manner approved by the Cambridge Psychology Research Ethics Committee, and were compensated for their participation at the rate of £7.50 per hour. For statistical analyses conducted on individual participant parameter estimates, we excluded outliers with a pre-defined criterion of a retrieval success (*pT*) or precision (*K*) estimate more than three standard deviations from the group mean. All participants were included in analyses conducted on parameters estimated from aggregate data (reported in Supplementary material).

### Data analysis approach

Retrieval error on each trial was calculated as the angular difference between participants’ response value and the target feature value (0 ± 180 degrees). To distinguish between different sources of memory errors (i.e., reduced retrieval success vs. reduced memory precision), a probabilistic mixture model was fitted to participants’ error data (Bays et al., 2009; Zhang & Luck, 2008) (code available at http://www.paulbays.com/code/JV10/index.php) (see Figure 1). In this model two sources of error contribute to participants’ performance: variability, that is, noise, in reporting the correct feature value when information about the target has been retrieved, and a proportion of trials where memory retrieval has failed and responses reflect random guessing. These two sources of error are modelled by two components: a von Mises distribution (circular equivalent of a Gaussian distribution) centred at a mean error of zero degrees from the target value, with a concentration *K*, and a circular uniform distribution with a probability *pU*. The concentration parameter, *K*, of the von Mises distribution captures variability in target retrieval (higher values reflect higher precision), and the probability of the uniform distribution, *pU,* reflects the likelihood of random guess responses, evenly distributed around the circular space. Of note, this model has previously been shown to best characterise participants’ long-term memory performance in an equivalent task (Richter, Cooper et al., 2016), and also fit the current data better than two alternative models for both younger and older participants (see Supplementary material).

**Figure 1.**
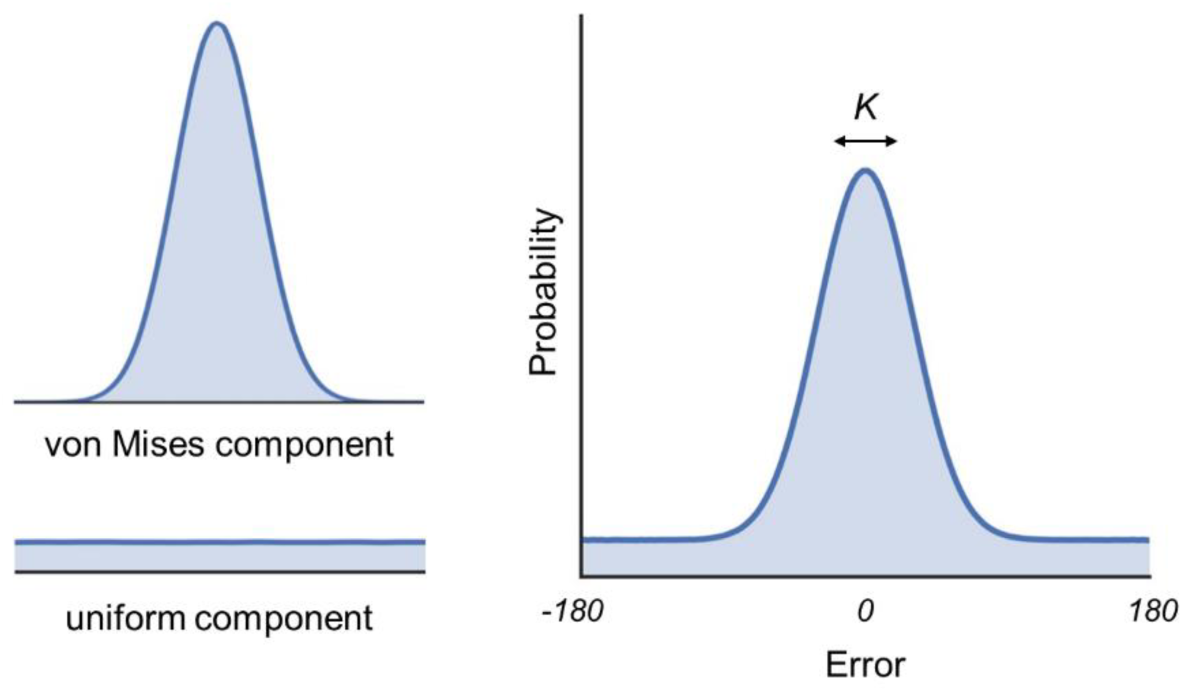
The probabilistic mixture model fit to participants’ retrieval error data consisted of a von Mises distribution (circular equivalent of a Gaussian distribution) centred at the target feature value, and a circular uniform distribution. Success of memory retrieval was defined as the probability of responses stemming from the target von Mises distribution (*pT*), and precision as the concentration (*K*) of the von Mises distribution.

The mixture model was fitted separately to data from each participant and task condition, yielding maximum likelihood estimates of the success (*pT*, probability of responses stemming from the target von Mises distribution, *pT* = 1 – *pU*) and precision (*K*, concentration of the von Mises distribution) of memory retrieval. Effects of group and task condition on the mean parameter estimates were assessed by t-tests and ANOVAs. We further validated the results obtained from modelling individual participants’ performance by modelling performance across all trials and participants in each age group. In all experiments, the results obtained by these two approaches converged. More detailed results of aggregate analyses are reported in the Supplementary material. Model fits were visualized with MATLAB MemToolbox (Suchow, Brady, Fougnie, & Alvarez, 2013; available at http://visionlab.github.io/MemToolbox/). Two-tailed *p*-values are reported for all analyses.

## Experiment 1

Experiment 1 employed a continuous location report task to examine whether age-related declines in episodic memory are attributable to reduced probability of successful memory retrieval, or to reduced precision of the retrieved memory representations. Selection of location as the tested feature in the first experiment was motivated by previous research demonstrating the retrieval of contextual, such as spatial, information as particularly vulnerable to age-related decline (Old & Naveh-Benjamin, 2008; Spencer & Raz, 1995). In the location memory task in Experiment 1, young and older participants encoded stimuli displays consisting of three everyday objects overlaid on a scene background. The location of each object on its associated background was pseudo-randomly selected from a circular space, and at retrieval, participants were asked to recreate the locations of studied objects by moving the object back to its original position as accurately as they could using a continuous response dial.

## Methods

### Participants

Twenty younger adults (19-23 years old), and 22 older adults (60-73 years old) participated in Experiment 1 (see Table 1). One older adult participant with a precision estimate > 3 *SDs* from the mean was excluded from the analyses conducted on the individual participant parameter estimates, leaving 20 younger and 21 older adults who contributed to the analyses. Older adults reported a higher number of years of formal education than younger adults, *t*(40) = 2.06, *p* = .046, *d* = 0.64. Moreover, older adults also had higher scores than younger adults on the SILVS (Zachary & Shipley, 1986), *t*(40) = 6.22, *p* < .001, *d* = 1.92, as typically observed in studies of cognitive ageing (Verhaeghen, 2003), indicating higher crystallized intelligence in the older group.

**Table 1.**
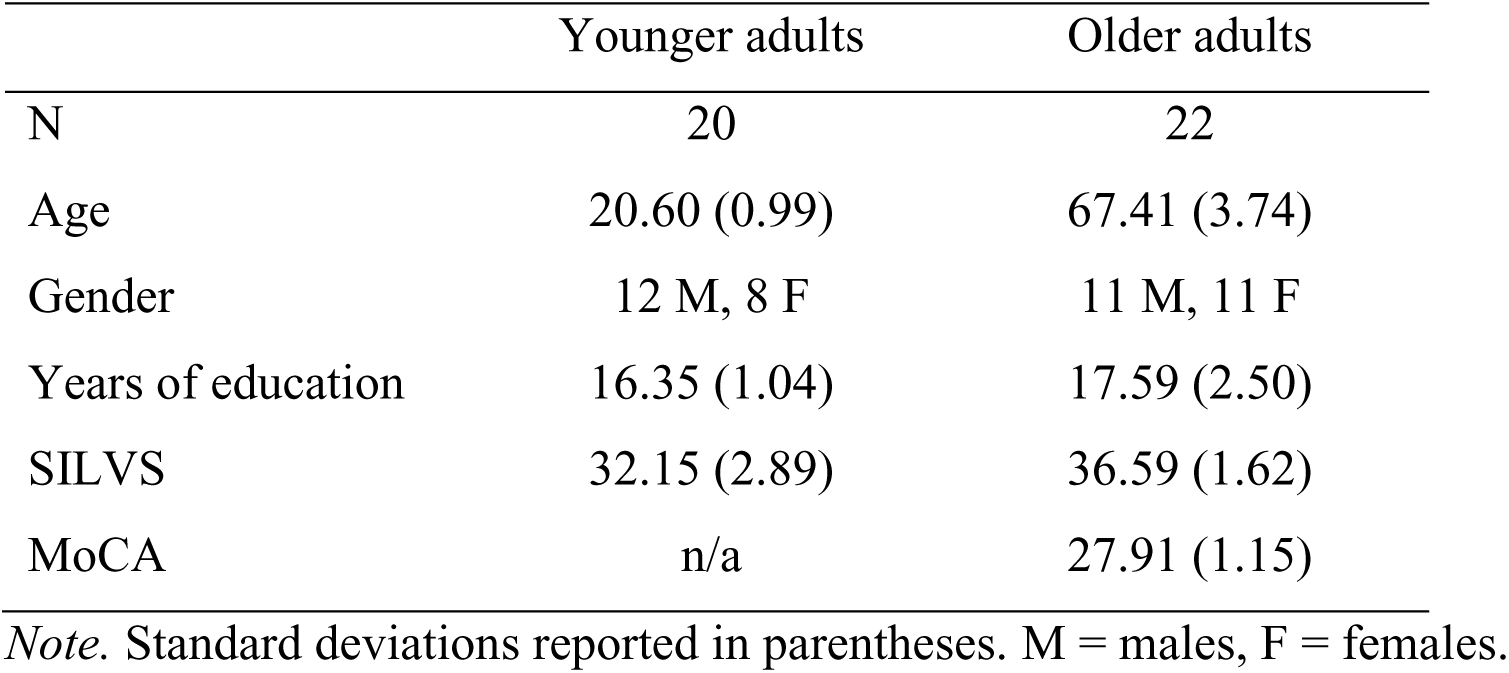
Participant demographic information in Experiment 1.

### Materials

The stimuli consisted of 180 images of distinct everyday objects, and 60 images of outdoor scenes. Object and scene images were obtained from existing stimuli sets (objects: Brady, Konkle, Alvarez, & Oliva, 2008; Konkle, Brady, Alvarez, & Oliva, 2010; scenes: Richter, Cooper et al., 2016) and Google image search. Three object images were randomly allocated to each scene image, forming a total of 60 trial-unique stimuli displays. The objects were each overlaid on the background scene in a location pseudo-randomly selected from a 360-degree circle with a radius of 247 pixels. A minimum distance of 62.04 degrees was enforced between the locations of any two objects on the same display, to ensure that the objects did not overlap. Displays were generated once, and all participants learned the same stimuli.

### Design and procedure

The location memory task consisted of 120 retrieval trials, divided into 5 study-test blocks (see Figure 2). In each study phase, participants viewed 12 stimuli displays for 9s each. The study phase was followed by a 30s delay, during which participants counted backwards by threes aloud, to prevent rehearsal of the studied stimuli. In the test phase, participants were first presented with a previously studied scene image with no objects overlaid on it for 9s, during which they were instructed to think about which objects had been associated with the given scene and where they had been located. Participants were then asked to sequentially reconstruct the locations of two out of three objects that had been associated with the scene as precisely as they could. Each object initially appeared in a random location on the associated background, along with a response dial. Participants were able to move the object clockwise and anti-clockwise around the 360-degree response dial by pressing the left and right arrow keys on the keyboard, and confirmed their answer by pressing the space bar. Response time was not limited to avoid disadvantaging the older adults; however participants were encouraged to try and respond within 15s. The passing of 15s was indicated by the central retrieval cue (“Location”) changing colour from white to red. Participants in both groups responded within the first 15 seconds on around 98% of trials. Participants completed 24 location retrieval trials in each block. Both encoding and retrieval trials were separated by a central fixation cross of 1s. The order of display presentation at study and test was randomised across participants. Which two out of the three studied objects per display were selected for location retrieval, and their test order, were randomized but kept constant across participants.

**Figure 2.**
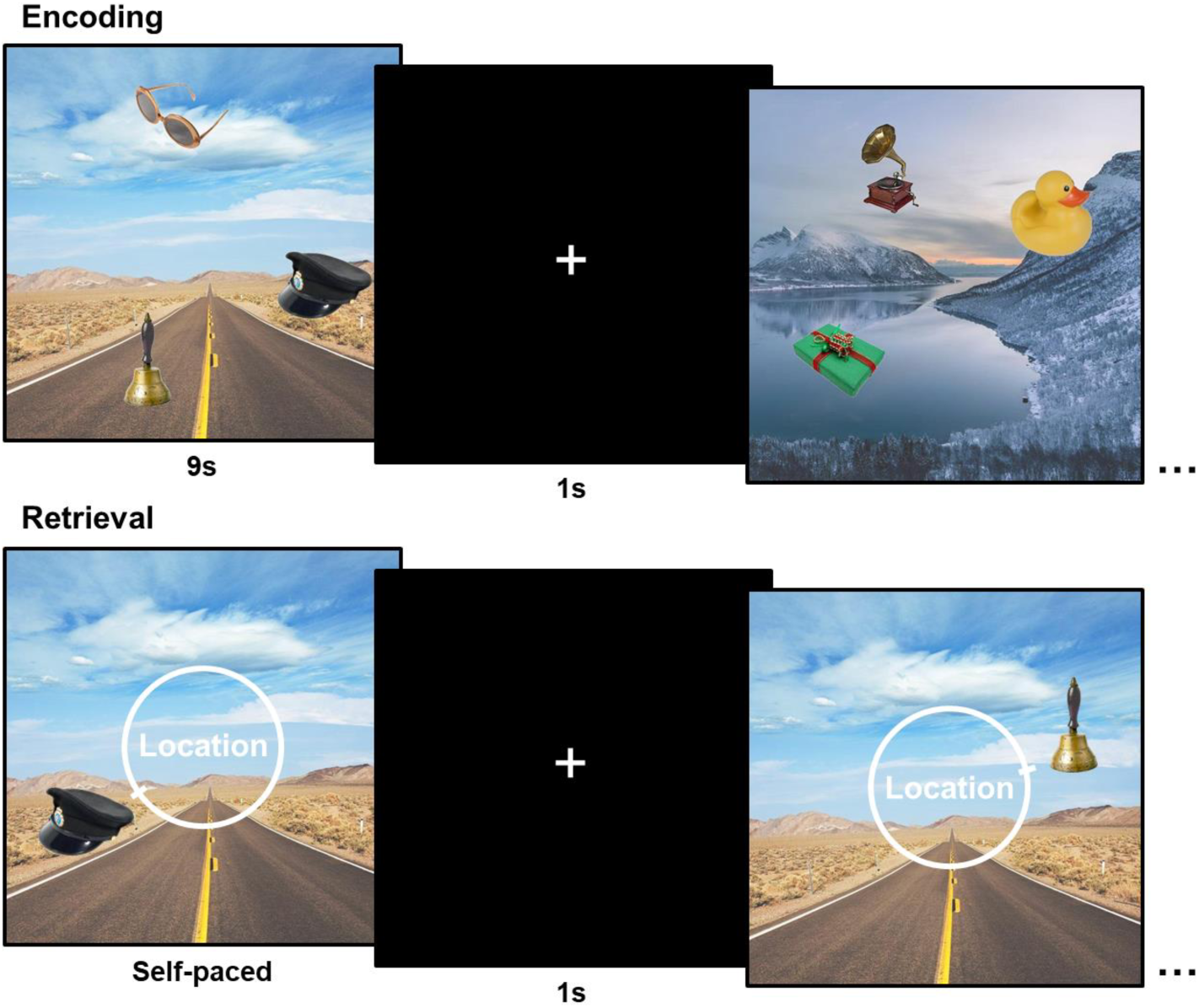
Example study and test trials in the location memory task in Experiment 1. Participants viewed stimuli displays (stimulus duration: 9s) consisting of three objects overlaid on a scene background, and later recreated the locations of two objects associated with each display, by moving the object around a 360-degree response dial via keypress. Retrieval error on each trial was calculated as the angular deviation between participants’ response value and the target location value (0 ± 180 degrees).

## Results

The distributions of participants’ retrieval errors (response value – target value) across the 120 retrieval trials in each age group are displayed in Figure 3, illustrating that on most trials participants recalled some information about the correct location with a variable degree of noise (proportion of errors centred around the target location), but on some trials memory retrieval failed leading to participants guessing a random location on the response dial (proportion of errors distributed uniformly across the circular space).

**Figure 3.**
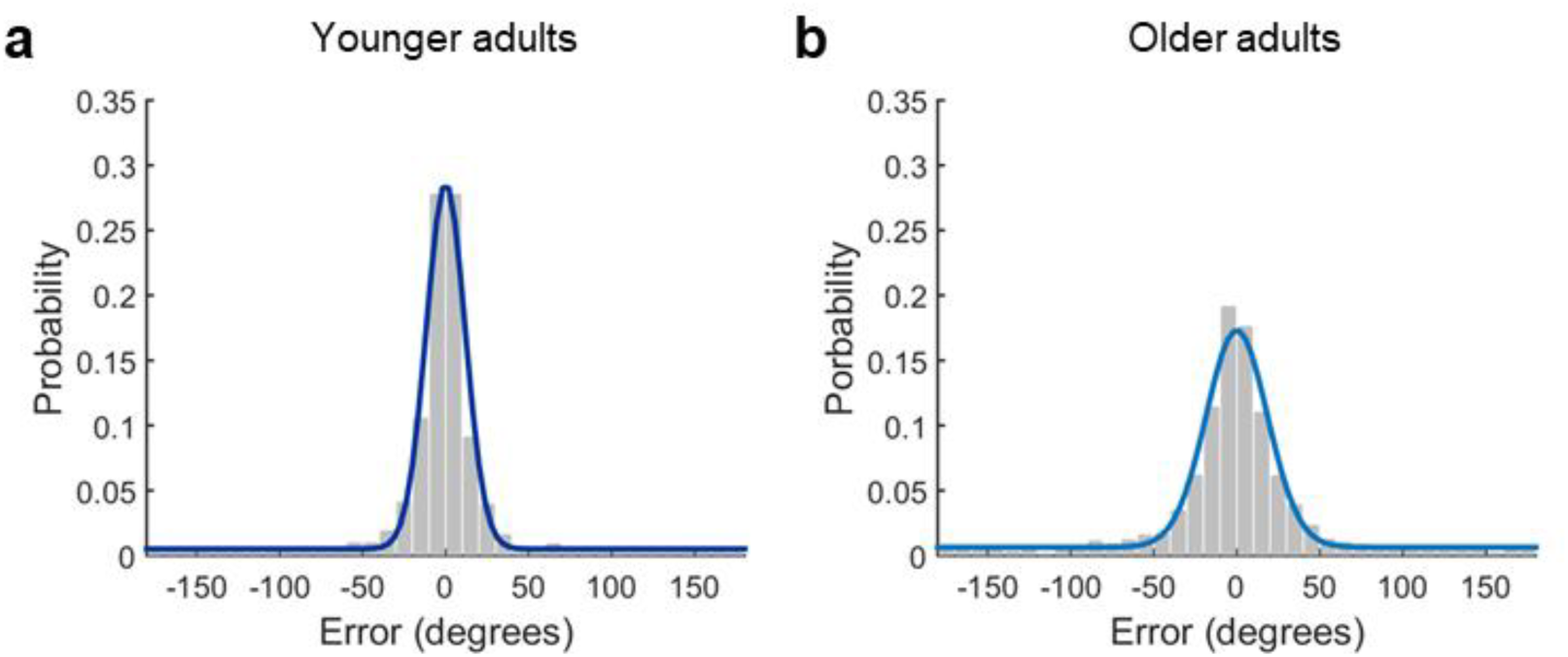
Distribution of retrieval errors (response feature value – target feature value) in the a) young and b) older adults. Coloured lines (dark blue: younger adults, light blue: older adults) indicate response probabilities predicted by the mixture model with target von Mises and circular uniform components (model fit to aggregate data in each group for visualization), illustrating similar retrieval success (equal height of the uniform components), but reduced memory precision in the older group (broader Gaussian component).

To quantify the success and precision of memory retrieval, we fitted a probabilistic mixture model (Bays et al., 2009; Zhang & Luck, 2008) to participants’ error data, yielding maximum likelihood estimates of the probability of successful memory retrieval (*pT*), and the precision of successful memory retrieval (*K*) for each participant. Examination of age differences in the model-derived estimates of retrieval success and precision indicated no significant differences in the mean probability of successful memory retrieval between the age groups, *t*(39) = 0.43, *p* = .669, *d* = 0.13 (see Figure 4a). However, the precision of memory retrieval was significantly reduced in the older group, *t*(39) = 4.96, *p* < .001, *d* = 1.55, indicating increased variability of target reports in older adults (see Figure 4b). To examine whether the observed age-related declines in retrieval precision were significantly greater than any age differences in retrieval success, we converted participants’ retrieval success and precision estimates to z-scores. A significant interaction between retrieval measure (retrieval success vs. precision) and age group (young vs. old), *F*(1, 39) = 6.00, *p* = .019, *partial η^2^* = 0.13, indicated disproportionate age-related declines in retrieval precision. The results of Experiment 1 therefore provide evidence for a selective deficit in the precision of location retrieval in the older group.

**Figure 4.**
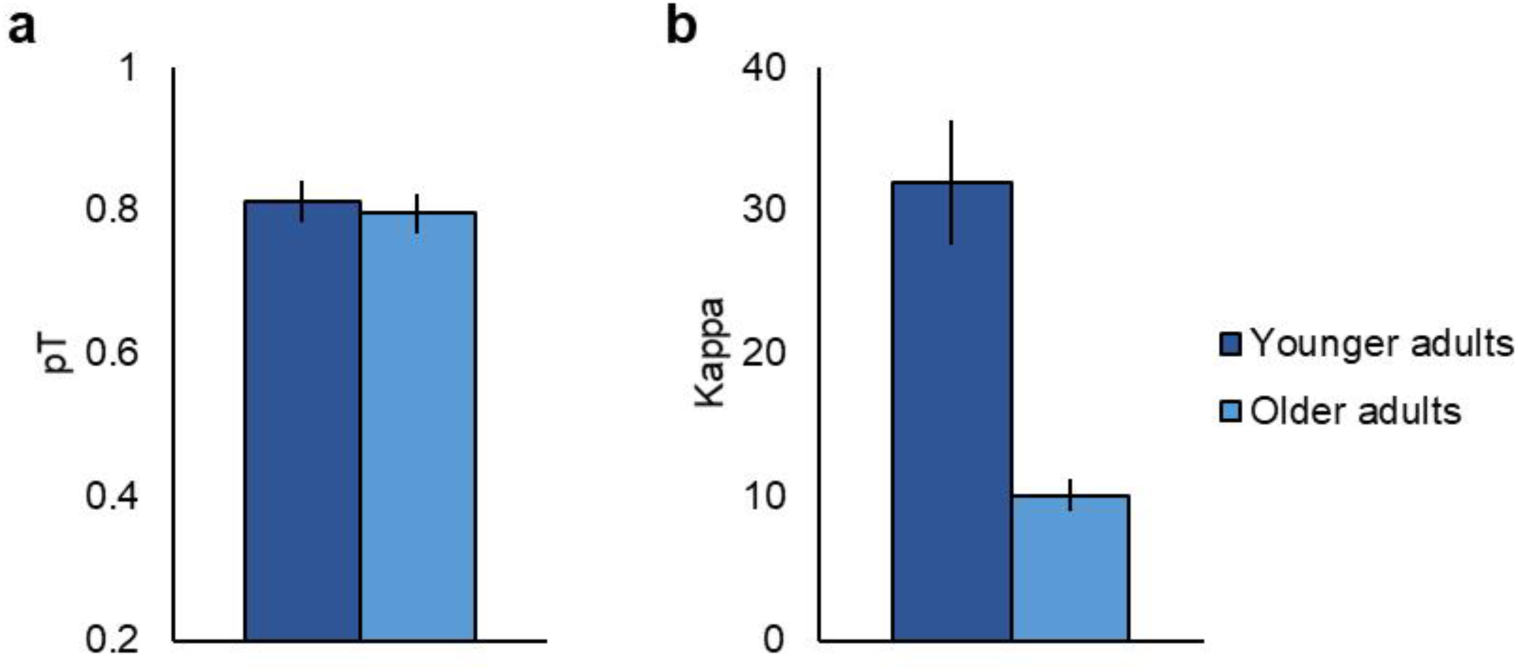
Mean a) retrieval success (*pT*) and b) retrieval precision (*K*) in each age group. Error bars display ± 1 standard error of the mean (SEM).

## Experiment 2

Following the finding of reduced precision of location memory retrieval in Experiment 1, we were next interested in exploring whether this deficit is specific to retrieval of contextual information, or evident across different types of information retained in LTM. Memory for objects themselves and the context in which they have been encountered have been argued to rely on dissociable neural circuits (Davachi, 2006; Ranganath & Ritchey, 2012), with retrieval of these two types of information also exhibiting a differential pattern of age-related decline (Old & Naveh-Benjamin, 2008; Spencer & Raz, 1995). In contrast to contextual retrieval, item-based memory has typically been considered more resilient to age-related degradation (Old & Naveh-Benjamin, 2008; Spencer & Raz, 1995). However, others have shown robust age-related declines for item memory also (e.g., Henson et al., 2016), in particular on tasks requiring a more detailed memory representation of the item to support accurate performance (Stark et al., 2013; Trelle, Henson, Green, & Simons, 2017), suggesting that the fidelity of item-based memory retrieval might also exhibit declines in older age. In the second experiment, we therefore examined the effects of ageing on the precision of both contextual and item-based feature retrieval, to test whether similar deficits in item memory can be detected using the continuous report paradigm. Participants encoded and retrieved stimuli displays consisting of three everyday objects varying in terms of their location (contextual), colour (item-based) and orientation (item-based) in circular spaces. At test, participants recreated the appearance of each feature using the continuous response dial. To investigate whether age-related decreases in the objective precision of memory retrieval were accompanied by reductions in the subjective quality of memories, we further asked participants to rate the subjective vividness of their memory retrieval for each display on a continuous scale.

## Methods

### Participants

Twenty-four younger (18-28 years old) and 24 older adults (62-79 years old) participated in Experiment 2 (see Table 2 for participant demographics). Six of the older adults had also participated in Experiment 1 (no overlap in task stimuli). Two younger adults and one older adult outlier (parameter estimates > 3 *SDs* from the mean) were excluded, leaving 22 younger adults and 23 older adults to contribute to the analyses based on individual parameter estimates. Older adults reported a significantly higher number of years of formal education than younger adults, *t*(46) = 2.17, *p* = .035, *d =* 0.63, and scored on average higher on the SILVS (Zachary & Shipley, 1986), *t*(45) = 4.55, *p* < .001, *d =* 1.33.

**Table 2.** Participant demographic information in Experiment 2.

|  | Younger adults | Older adults |
| --- | --- | --- |
| N | 24 | 24 |
| Age | 21.00 (2.40) | 71.83 (4.57) |
| Gender | 12 M, 12 F | 12 M, 12 F |
| Years of education | 16.04 (1.40) | 17.54 (3.08) |
| SILVS<sup>a</sup> | 33.75 (3.53) | 37.48 (1.75) |
| MoCA | n/a | 28.13 (1.23) |

*Note.* Standard deviations reported in parentheses.
<sup>a</sup> SILVS score missing for one older adult due to experimenter error.

### Materials

Stimuli for the continuous report task in Experiment 2 consisted of 120 images of distinct everyday objects and 40 images of textured backgrounds. The object images were obtained from an existing stimuli set (Brady et al., 2013, available at http://timbrady.org/stimuli/ColorRotationStimuli.zip), and the background images from Google Image Search (no overlap with Experiment 1). The stimuli were randomly allocated to form a total of 40 trial-unique study displays each consisting of three objects overlaid on a texture background. In contrast to Experiment 1, in Experiment 2 the objects on each display varied along three perceptual features: location, colour and orientation. Values for each of these features were pseudo-randomly drawn from a circular space (0-360 degrees) with the constraint of a minimum distance of 62.04 degrees between two features of the same type on each display. As in Experiment 1, this minimum distance was required to create non-overlapping object locations, and for consistency also applied to the other two feature dimensions. All participants learned the same displays.

### Design and procedure

The continuous report task consisted of 10 study-test blocks (see Figure 5). In each study phase, participants sequentially viewed four stimuli displays (stimulus duration: 12s), after which a 10s delay followed. In the test phase, participants were first asked to rate the vividness of their memory for each display, and to base this vividness judgement on how vividly they could recall the appearance of all of the three objects associated with that display. Participants were presented with the background image only, along with a question “How vividly do you remember this display?” in the centre of the image. After 2s delay a response scale was added and participants could indicate the vividness of their memory by moving a slider on a 100-point continuous scale (0 = “not vivid”, 100 = “very vivid”). After the vividness rating, participants sequentially reconstructed the features (location, colour, and orientation) of two out of three objects on each display. For feature retrieval, the test object initially appeared in a randomly allocated location, colour and orientation on the associated background along with the response dial. A central cue noted the feature being tested (“Location”, “Colour”, or “Orientation”), and after responding to one of the feature questions, participants’ reconstruction of that feature’s appearance remained unchanged for the following feature questions for the same object. As in Experiment 1, the test phase was self-paced, but participants were encouraged to respond within 15s, with the retrieval prompt on the screen (i.e., the vividness question or the feature label) changing colour from white to red after 15s. The study and test trials were separated by a fixation cross of jittered duration (400ms to 2500ms, mean: 1025ms).

**Figure 5.**
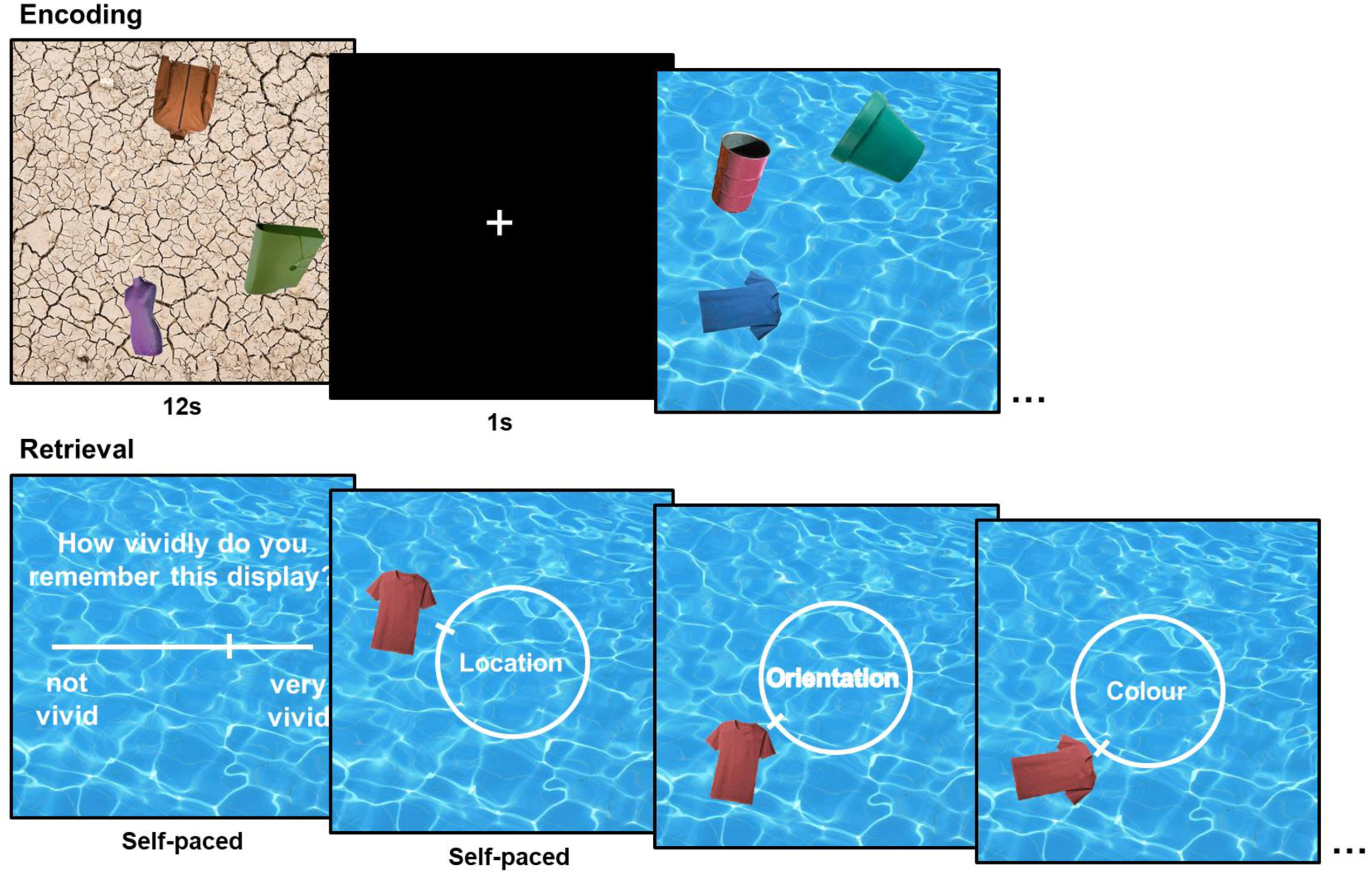
In Experiment 2, participants studied stimuli displays consisting of three objects varying along three features: location, colour and orientation (stimulus duration: 12s). For each display, participants first rated the vividness of their memory retrieval, and then recreated the features of two out of three objects on each display, using the continuous response dial.

Participants completed 40 vividness trials, and 240 feature retrieval trials (80 per feature) in total. The order of display presentation at study and test was randomized across participants. Selection of two objects from each display for feature retrieval and their test order was randomized but kept constant across participants. The order of feature questions for each object was pseudo-randomised across participants with the constraint of no individual feature tested more than 4 consecutive times in the same sequential position (i.e., first, second, or third).

## Results

Distributions of retrieval errors in each feature condition and age group in Experiment 2 are displayed in Figure 6. Analysis of mean parameter estimates indicated that age differences in retrieval success varied significantly across the three feature conditions, *F*(2, 86) = 5.03, *p* = .009, *partial η^2^* = 0.11 (see Figure 7a). Whereas no significant age differences in retrieval success were observed in the location, *t*(43) = 0.13, *p* = .901, *d* = 0.04, and colour, *t*(43) = 1.34, *p* = .186, *d* = 0.40, conditions, older adults exhibited significantly lower probability of successful memory retrieval than younger adults in the orientation condition, *t*(43) = 2.45, *p* = .018, *d* = 0.73. The orientation condition further had the lowest retrieval success out of the three feature conditions across both age groups (lower retrieval success than colour, *t*(44) = 6.47, *p* < .001, *d* = 1.81, and location, *t*(44) = 12.14, *p* < .001, *d* = .96), indicating that the only significant age differences in retrieval success were observed for the condition that also resulted in the poorest overall performance. No significant main effect of age group was observed for retrieval success, *F*(1, 43) = 2.29, *p* = .138, *partial η^2^* = 0.05.

**Figure 6.**
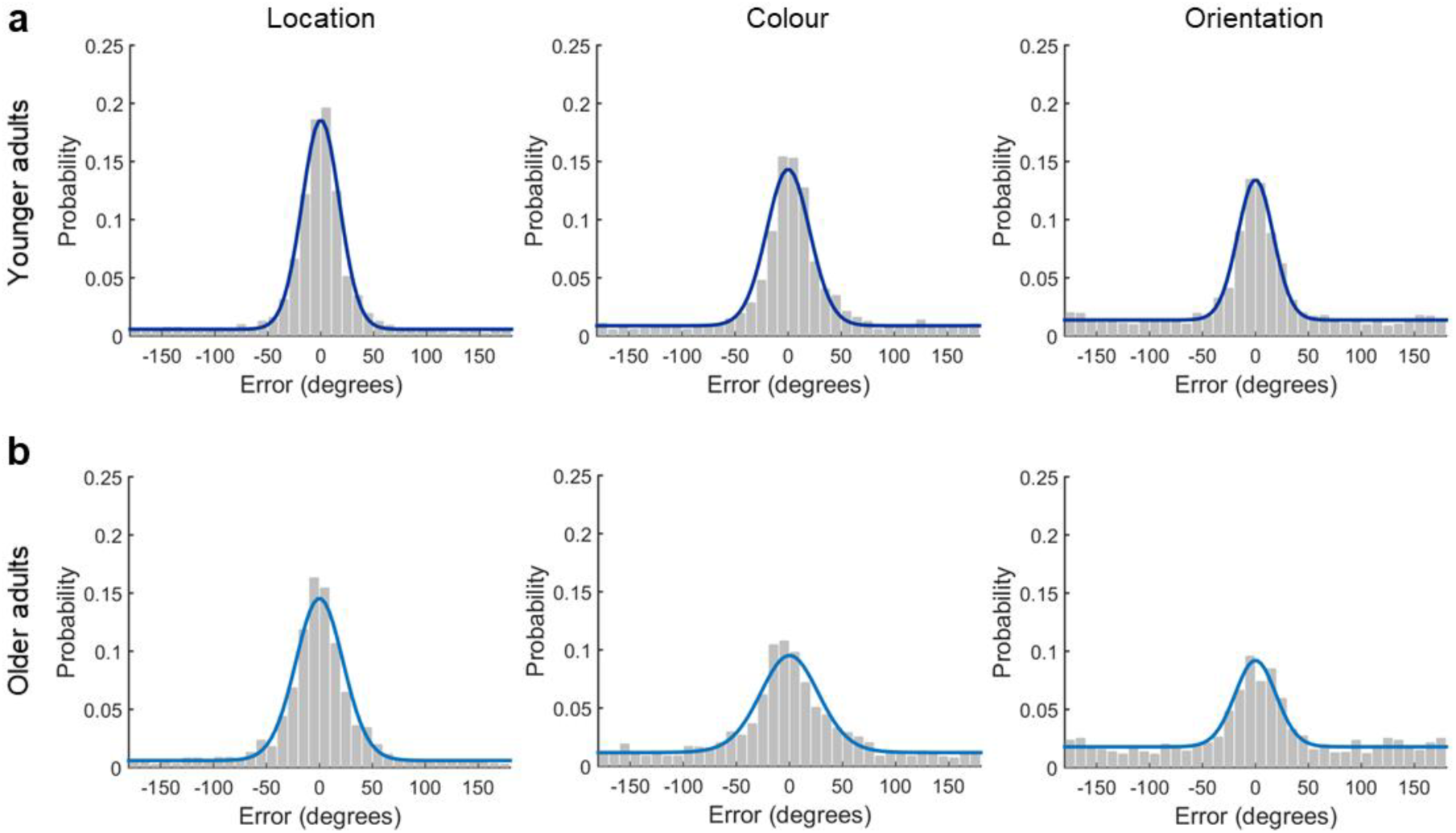
Distribution of retrieval errors in each feature condition in the a) younger and b) older adults. Coloured lines (dark blue: younger adults, light blue: older adults) illustrate response probabilities predicted by the mixture model (model fit to aggregate data for visualization).

**Figure 7.**
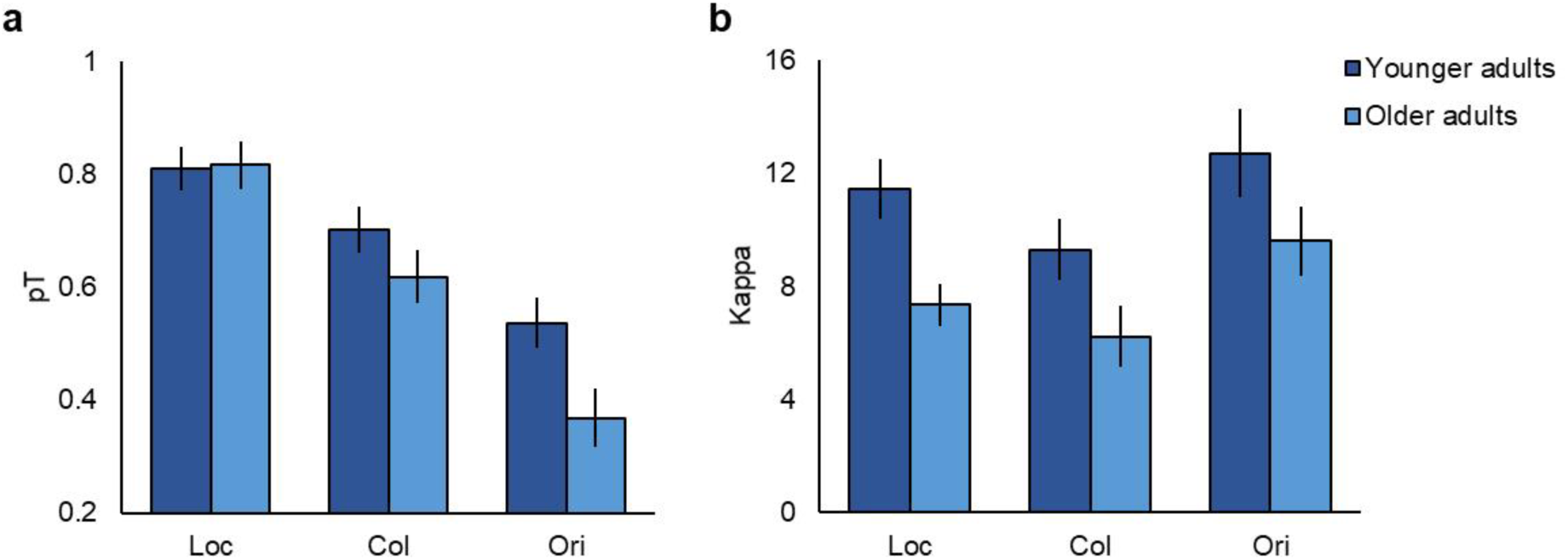
Mean a) retrieval success (*pT*) and b) retrieval precision (*Κ*) in each age group and feature condition (Loc = location, Col = colour, Ori = orientation). Error bars display ± 1 SEM.

In contrast, for retrieval precision, we observed a significant main effect of age group, *F*(1, 43) = 11.54, *p* = .001, *partial η^2^* = 0.21, indicating reduced precision of memory retrieval in the older group (see Figure 7b). Age differences in retrieval precision did not vary significantly across the feature conditions, *F*(2, 86) = 0.14, *p* = .872, *partial η^2^* = 0.00, indicating age-related loss of mnemonic precision across different types of information retained in LTM.

Comparing the magnitude of age differences in retrieval success and precision for each feature condition, we found evidence for a disproportionate deficit in retrieval precision in the location condition, *F*(1, 43) = 5.52, *p* = .023, *partial η^2^* = 0.11, but not in the colour, *F*(1, 43) = 0.17, *p* = .679, *partial η^2^* = 0.00, or orientation, *F*(1, 43) = 0.33, *p* = .570, *partial η^2^* = 0.01, conditions (estimates on each measure z-scored).

Furthermore, the mean subjective ratings of memory vividness did not significantly differ between the age groups, *t*(43) = 0.71, *p* = .485, *d* = 0.21, with both age groups rating their memory retrieval as moderately vivid (younger: *M*: 45.09, *SD*: 13.12, older: *M*: 49.65, *SD*: 27.42, on a scale 0-100).

## Discussion

In Experiment 2, we assessed the fidelity of participants’ long-term memory retrieval for both contextual (location) and item-based (colour and orientation) information. Consistent with results from the location memory task in Experiment 1, we here observed significant age-related declines in the precision of episodic memory across the three features tested, indicating loss of fidelity of both contextual and item-specific memory representations in older age.

In contrast to Experiment 1, in the present experiment we also observed significantly reduced probability of successful memory retrieval in the older group in the orientation condition. The orientation condition resulted in lower retrieval success than the other two feature conditions in both age groups, potentially suggesting that age differences in retrieval success might emerge with increasing task difficulty (Reuter-Lorenz & Cappell, 2008). Furthermore, despite reductions in the objective precision of memories, older adults did not display decreases in the subjective vividness of their memory retrieval, consistent with previous reports (Johnson, Kuhl, Mitchell, Ankudowich, & Durbin, 2015; St-Laurent et al., 2014).

## Experiment 3

Experiments 1 and 2 both revealed age-related deficits in the precision of episodic memory retrieval, however it is unclear how specific these decreases are to long-term memory, or whether they might at least partly reflect age-related declines in the fidelity of information processed at the level of perception or working memory. As the precision of mnemonic representations is constrained by the fidelity of sensory inputs (Ma, Husain, & Bays, 2014), age-related reductions in perceptual processing might contribute to the loss of memory precision observed in the current experiments, consistent with the information degradation hypothesis of age-related cognitive decline (Monge & Madden, 2016). Alternatively, or additionally, age-related limitations in episodic memory precision might arise from decreases in the precision of WM, documented in previous studies (Peich et al., 2013; Pertzov et al., 2015). Previous work in younger adults has proposed LTM and WM to exhibit similar constraints on representational fidelity (Brady et al., 2013), supporting the hypothesised link between age-related decreases in the precision of WM and LTM.

The aim of the third experiment was therefore to examine whether age-related declines in the precision of episodic memory could be partially, or fully, explained by declines in the fidelity of perceptual and/or working memory representations. In Experiment 3, participants completed perceptual, WM, and LTM versions of the continuous report task for object colour. Colour was chosen as the tested feature in this experiment as previous research in younger adults employing a similar task has found colour to be a sufficiently sensitive feature for investigating the fidelity of all three cognitive functions: perception, WM and LTM (Brady et al., 2013).

## Methods

### Participants

Twenty-six younger (18-30 years old), and 24 older adults (60-82 years old) took part in Experiment 3. Two younger adults were excluded from the experiment prior to data analysis, one due to a counterbalancing error leading to the participant completing the same tasks twice, and one due to failure to attend the second study session (see Table 3 for participant demographics for the remaining participants, 24 younger, and 24 older adults). Furthermore, two younger and two older adult outliers (parameter estimates > 3 *SDs* from the group mean) were excluded from the analyses based on individual participant parameter estimates, leaving 22 younger and 22 older adults to contribute to the analyses. Similar to previous experiments, older adults reported a significantly higher number of years of formal education than younger adults, *t*(46) = 2.19, *p* = .034, *d* = 0.63, and scored significantly higher on the SILVS (Nasreddine et al., 2005), *t*(46) = 6.58, *p* < .001, *d* = 1.90.

**Table 3.**
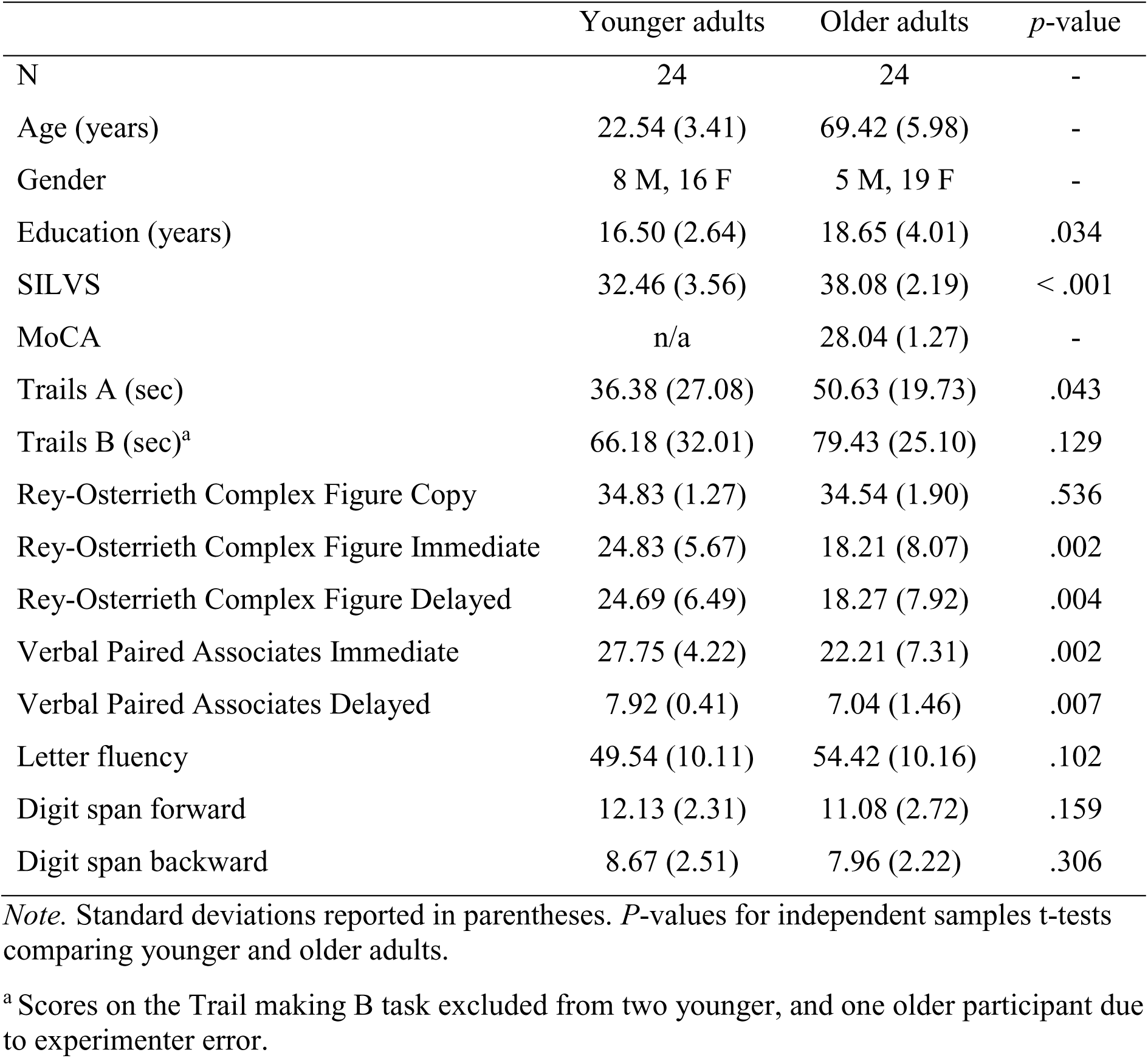
Participant demographic and neuropsychological test data in Experiment 3.

|  | Younger adults | Older adults | *p*-value |
| --- | --- | --- | --- |
| N | 24 | 24 | - |
| Age (years) | 22.54 (3.41) | 69.42 (5.98) | - |
| Gender | 8 M, 16 F | 5 M, 19 F | - |
| Education (years) | 16.50 (2.64) | 18.65 (4.01) | .034 |
| SILVS | 32.46 (3.56) | 38.08 (2.19) | < .001 |
| MoCA | n/a | 28.04 (1.27) | - |
| Trails A (sec) | 36.38 (27.08) | 50.63 (19.73) | .043 |
| Trails B (sec)<sup>a</sup> | 66.18 (32.01) | 79.43 (25.10) | .129 |
| Rey-Osterrieth Complex Figure Copy | 34.83 (1.27) | 34.54 (1.90) | .536 |
| Rey-Osterrieth Complex Figure Immediate | 24.83 (5.67) | 18.21 (8.07) | .002 |
| Rey-Osterrieth Complex Figure Delayed | 24.69 (6.49) | 18.27 (7.92) | .004 |
| Verbal Paired Associates Immediate | 27.75 (4.22) | 22.21 (7.31) | .002 |
| Verbal Paired Associates Delayed | 7.92 (0.41) | 7.04 (1.46) | .007 |
| Letter fluency | 49.54 (10.11) | 54.42 (10.16) | .102 |
| Digit span forward | 12.13 (2.31) | 11.08 (2.72) | .159 |
| Digit span backward | 8.67 (2.51) | 7.96 (2.22) | .306 |

*Note.* Standard deviations reported in parentheses. *P*-values for independent samples *t*-tests comparing younger and older adults.
<sup>a</sup> Scores on the Trail making B task excluded from two younger, and one older participant due to experimenter error.

### Materials

Stimuli for all tasks consisted of 540 everyday objects (Brady et al., 2008; Brady et al., 2013), including the object stimuli from Experiment 2 (no overlap in participants). Objects that were not readily colour-rotated were initially converted to the same hue of red as the Brady et al. (2013) colour-rotated object stimuli. Object images were randomly allocated to each task type, with 120 objects allocated to the LTM task, 360 to the WM task, and 60 to the perceptual task. In the LTM and WM tasks, stimuli displays consisted of three distinct objects overlaid on a grey background. To keep the amount of visual input consistent across tasks, stimuli displays in the perception task also comprised three objects overlaid on a grey background. However, as this task involved no demands on memory, three versions of the same object were used. The colour and location of the objects in each display were pseudo-randomly chosen from circular parameter spaces with the minimum constraint of 62.04 degrees between two feature values of the same type. A total of 40 unique stimuli displays were created for the LTM task, 120 for the WM task, and 60 for the perception task. All participants viewed the same displays.

### Design and procedure

Participants attended two testing sessions, with a minimum one week delay between the sessions (delay for younger adults *M:* 11.89 days, *SD:* 7.02, older adults *M:* 11.46 days, *SD:* 7.26, no significant difference between the groups, *t*(46) = 0.20, *p* = .841, *d* = 0.06). In addition to the three colour report tasks, participants completed a battery of standard neuropsychological tasks including measures of verbal (Verbal Paired Associates, WMS-III) (Wechsler, 1997b) and non-verbal memory (Rey-Osterrieth Complex Figure test) (Osterrieth, 1944), executive function (Verbal fluency, Trails A & B) (Delis, Kaplan, & Kramer, 2001), and working memory (Digit span forward and backward, WAIS-III) (Wechsler, 1997a). Participants’ performance on the neuropsychological tasks is presented in Table 2. The assignment of the colour report tasks and neuropsychological tests to each testing session was counterbalanced across participants, with the memory versions of the task completed in separate sessions.

### Colour report tasks

During the test phase of each of the continuous report tasks, the target object initially appeared in a randomly allocated colour but in its studied location and orientation (memory for location or orientation not tested in Experiment 3). As in the previous experiments, response time was not limited in any of the tasks, but participants were encouraged to respond within 15s, after which the retrieval cue (“Colour”) changed colour from white to red. Trials in each task were separated by a 1s central fixation cross. The order of displays at study and test was randomised across participants. The order of the objects to test per display in the LTM task (3 objects tested for each display), and the selection of objects to test per display in the WM and perceptual tasks (one object tested for each display) was randomised, but kept constant across participants.

The LTM task consisted of 120 colour retrieval trials, divided into 8 study-test blocks (see Figure 8). In each study phase, participants sequentially viewed five stimuli displays (stimulus duration: 9s). The study phase was followed by a 30s delay filled with counting backwards by threes aloud, to prevent rehearsal of the studied stimuli and to ensure that the task relied on long-term memory. In the test phase, participants recreated the colours of all the objects studied in the preceding block (15 retrieval trials per block).

**Figure 8.**
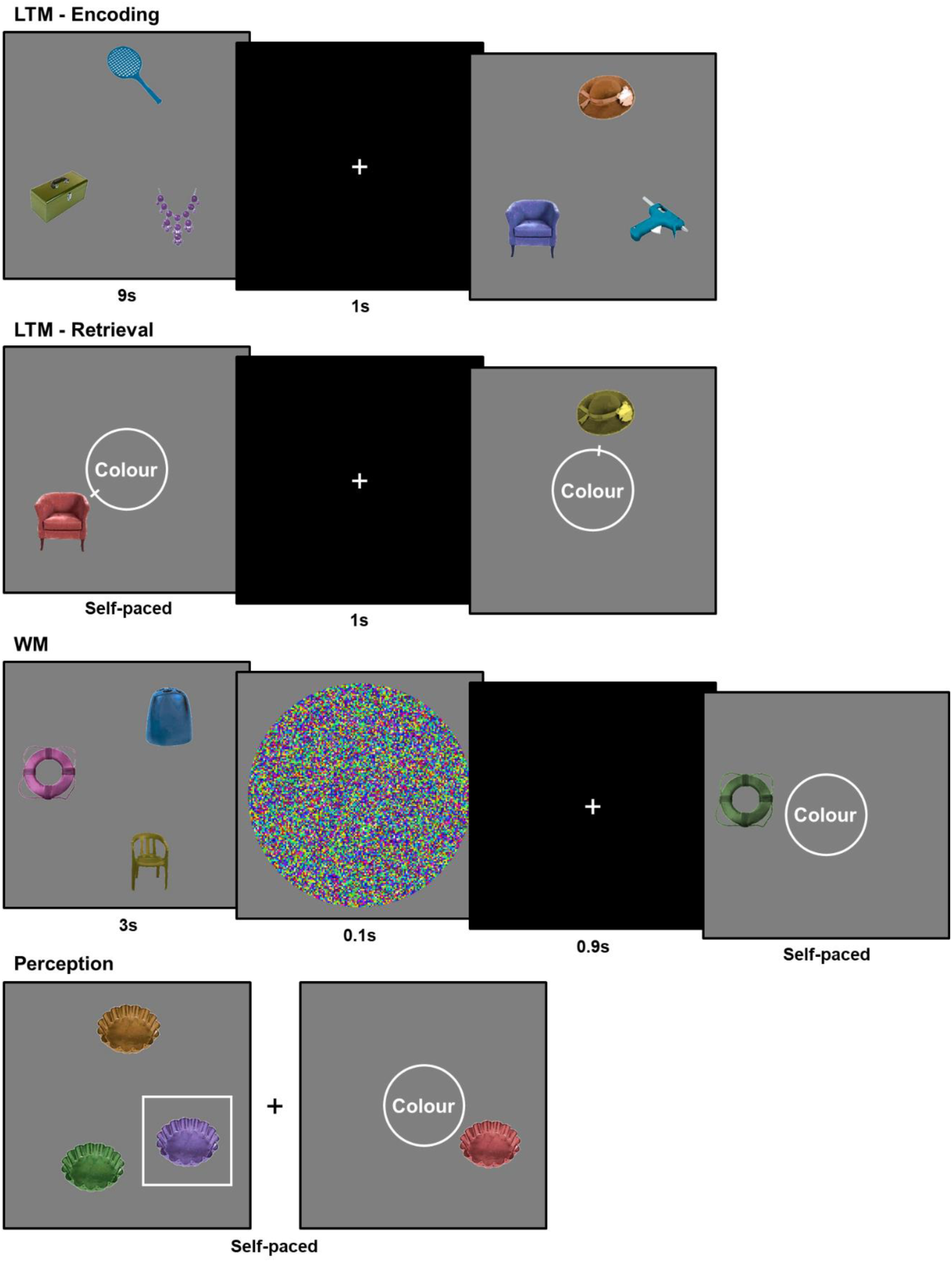
Example trials in each of the three colour report tasks. In the LTM task participants studied five stimuli displays in a row (9s each), before retrieving the colours of all objects after a 30s delay. In the WM task, participants studied one stimuli display at a time (3s each), and retrieved the colour of one object after 1s delay. In the perception task, participants matched the colour of one object per display while the stimuli display was simultaneously in view.

In the WM task, participants also completed 120 colour retrieval trials in total, divided into 8 blocks of 15 trials each (see Figure 8). In this task, participants studied only one stimuli display at a time (stimulus duration: 3s). To prevent reliance on sensory memory, display presentation was followed by presentation of a coloured mask image for 100ms, followed by a 900ms central fixation cross. After the total delay of 1s, participants reconstructed the colour of one of the objects from the preceding display. Participants were only tested on one object per display to ensure consistent demands on working memory.

The perceptual task included 60 trials, divided into two blocks of 30 trials each (see Figure 8). On each trial, participants saw two displays side-by-side on the screen. One of the displays had three versions of the same object presented in different colours. The other display had only one object, the colour of which participants were able to adjust with the response dial. The participants’ task was to match the colour of the test object to the colour of an object in the same relative location on the other display, surrounded by a white square. As the display was simultaneously in view, this task placed no demands on memory. The side of presentation of the display and test object on each trial (left vs. right) was randomised across participants.

## Results

As in both of the memory tasks, errors in the perception task were calculated as the angular deviation between participants’ response value and the target colour value (0±180 degrees). Distribution of errors in each task and age group are displayed in Figure 9.

**Figure 9.**
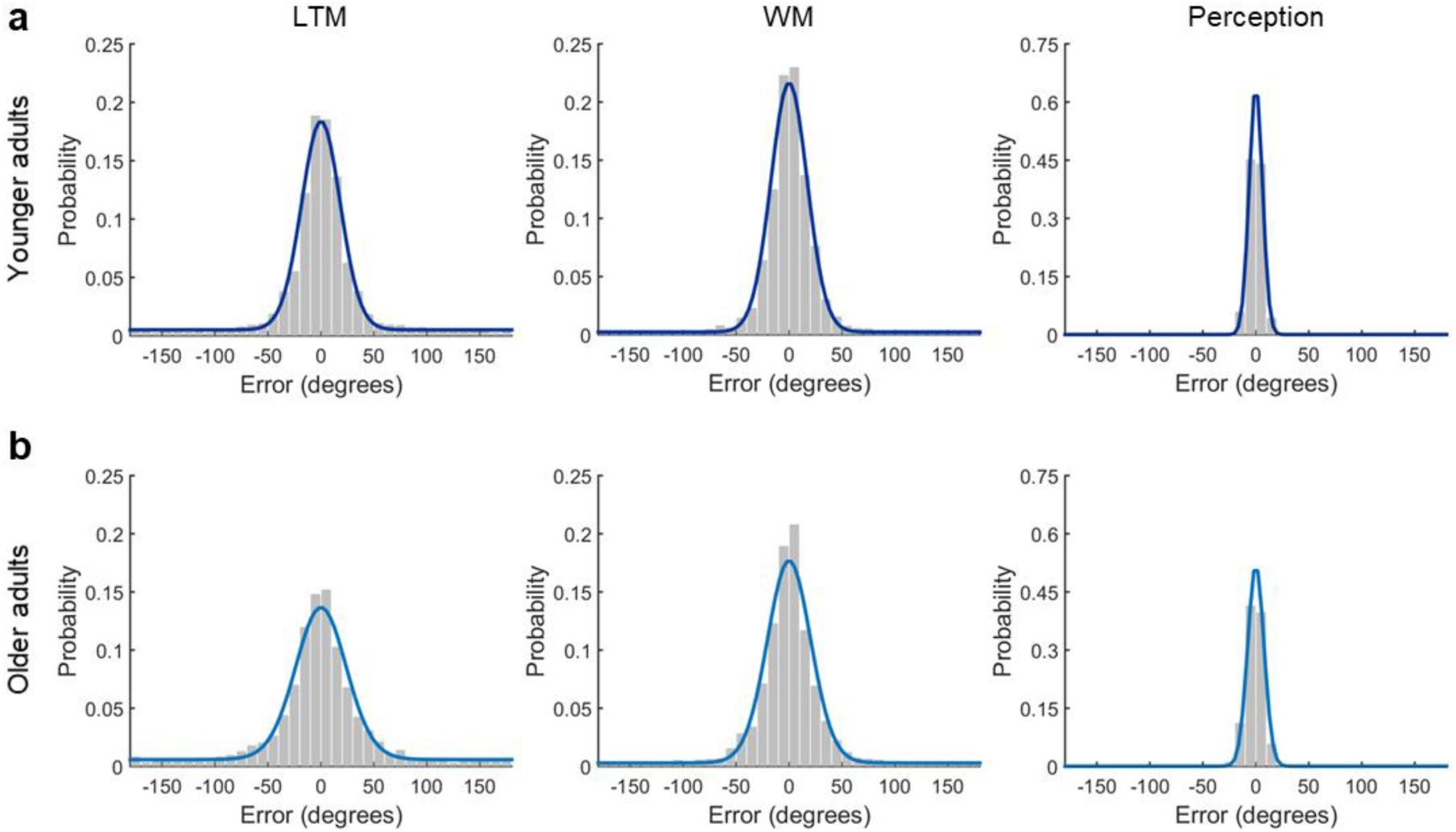
Distribution of errors in the LTM, WM and perception colour report tasks in a) younger and b) older adults. Coloured lines (dark blue: younger adults, light blue: older adults) illustrate response probabilities predicted by the mixture model (model fit to aggregate data for visualization). Note the different scaling of the y-axes for the perception task.

Focusing first on long-term memory, consistent with Experiments 1 (location condition) and 2 (location and colour conditions), comparison of mean parameter estimates indicated no significant differences in the probability of successful long-term memory retrieval between the age groups, *t*(42) = 0.76, *p* = .454, *d* = 0.23, but a significant decline in memory precision in the older group, *t*(42) = 4.12, *p* < .001, *d* = 1.24 (see Figures 10a and 10b). The deficit in LTM precision was disproportionate to any age differences in LTM retrieval success, *F*(1, 42) = 4.26, *p* = .045, *partial η^2^* = 0.09 (retrieval success and precision estimates z-scored). Similarly, in working memory, we observed no significant age differences in the probability of successful memory retrieval, *t*(42) = 0.80, *p* = .428, *d* = 0.24, but the older group displayed a significant reduction in memory precision, *t*(42) = 3.12, *p* = .003, *d* = 0.94 (see Figures 10a and 10b). However, the evidence for a disproportionate deficit in memory precision in the WM task was not significant, *F*(1, 42) = 2.15, *p* = .150, *partial η^2^* = 0.05 (estimates z-scored). Lastly, the age groups did not differ significantly in terms of the probability of reporting the correct target colour in the perception task, *t*(42) = 1.70, *p* = .097, *d* = 0.51 (see Figure 10a), during which the stimuli display was simultaneously in view as the participants selected their response. However, even in the perceptual task, older adults were significantly less precise at matching the colour of the objects than younger adults, *t*(42) = 3.40, *p* = .001, *d* = 1.03 (see Figure 10c). The evidence for a disproportionate deficit in precision in the perceptual task was not significant, *F*(1, 42) = 1.33, *p* = .256, *partial η^2^* = 0.03 (estimates z-scored).

**Figure 10.**
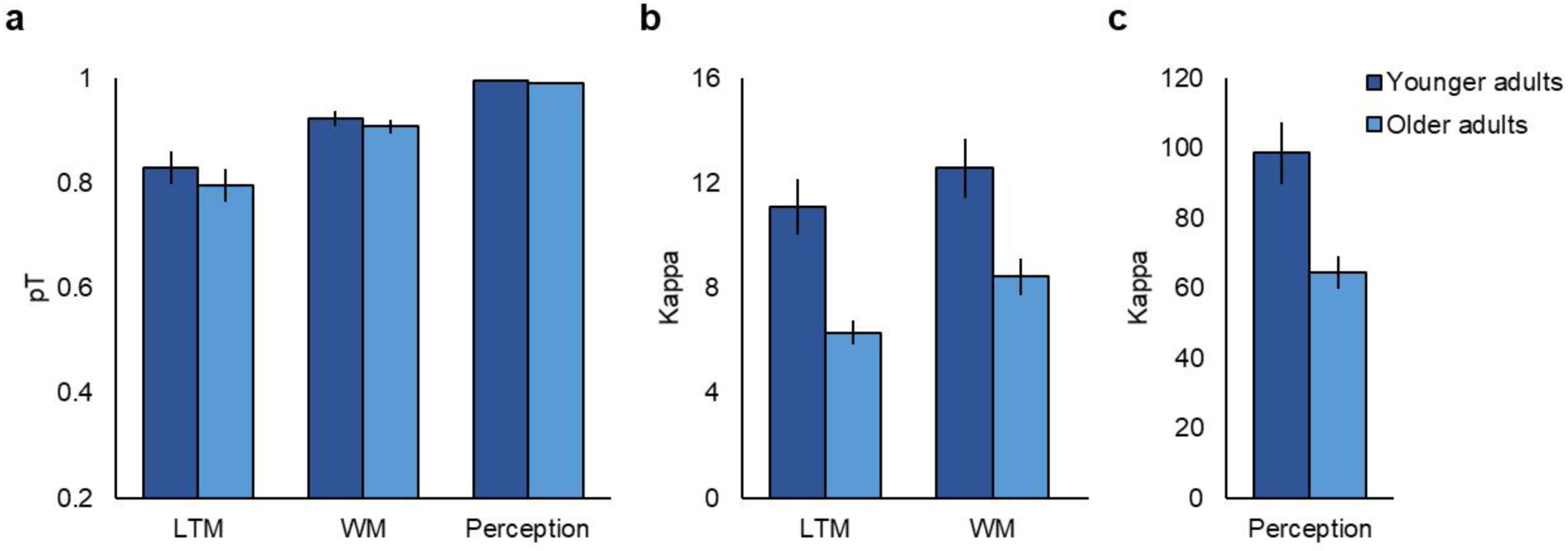
Mean a) probability of target reports (*pT*) and b), c) precision (*Κ*) in each age group. Note that precision in the perceptual task is plotted separately due to higher *Κ* values in this task. Error bars display ± 1 SEM.

To investigate whether variation in perception or working memory predicted long-term memory performance in each of the age groups, we used linear regression (see Figure 11). In younger adults, a model including both perceptual and WM precision as predictor variables did not significantly predict the precision of LTM retrieval, *R^2^* = .12, *F*(2, 21) = 1.24, *p* = .311, nor did either of these two variables alone (perception: *R^2^* = .11, *F*(1, 21) = 2.54, *p* = .126; WM: *R^2^* = .01, *F*(1, 21) = 0.14, *p* = .717). In contrast, in the older group, a model including both perceptual and WM precision was a significant predictor of LTM precision, *R^2^* = .36, *F*(2, 21) = 5.45, *p* = .014. However, this result was driven by a significant effect of WM precision on LTM precision, *R^2^* = .35, *F*(1, 21) = 10.67, *p* = .004, whereas perceptual precision alone did not significantly predict LTM precision in the older group, *R^2^* = .12, *F*(1, 21) = 2.68, *p* = .117. Furthermore, perceptual precision did not significantly predict WM precision in either younger, *R^2^* =.15, *F*(1, 21) = 3.60, *p* = .072, or older adults, *R^2^* = .15, *F*(1, 21) = 3.41, *p* = .080, and retrieval success in the WM task did not significantly predict LTM retrieval success (young: *R^2^* = .16, *F*(1, 21) = 3.68, *p* = .069, old: *R^2^* = .00, *F*(1, 21) = 0.05, *p* = .825), or LTM precision (young: *R^2^* = .01, *F*(1, 21) = 0.15, *p* = 705, old: *R^2^* = .01, *F*(1, 21) = 0.16, *p* = .698) in either age group. Note that we did not examine the relationship between probability of successful target reports in the perceptual task and other performance measures due to lack of variability on this measure in both age groups (younger adults *M*: 1.00, *SD*: 0.01, older adults *M*: 0.99, *SD*: 0.01).

**Figure 11.**
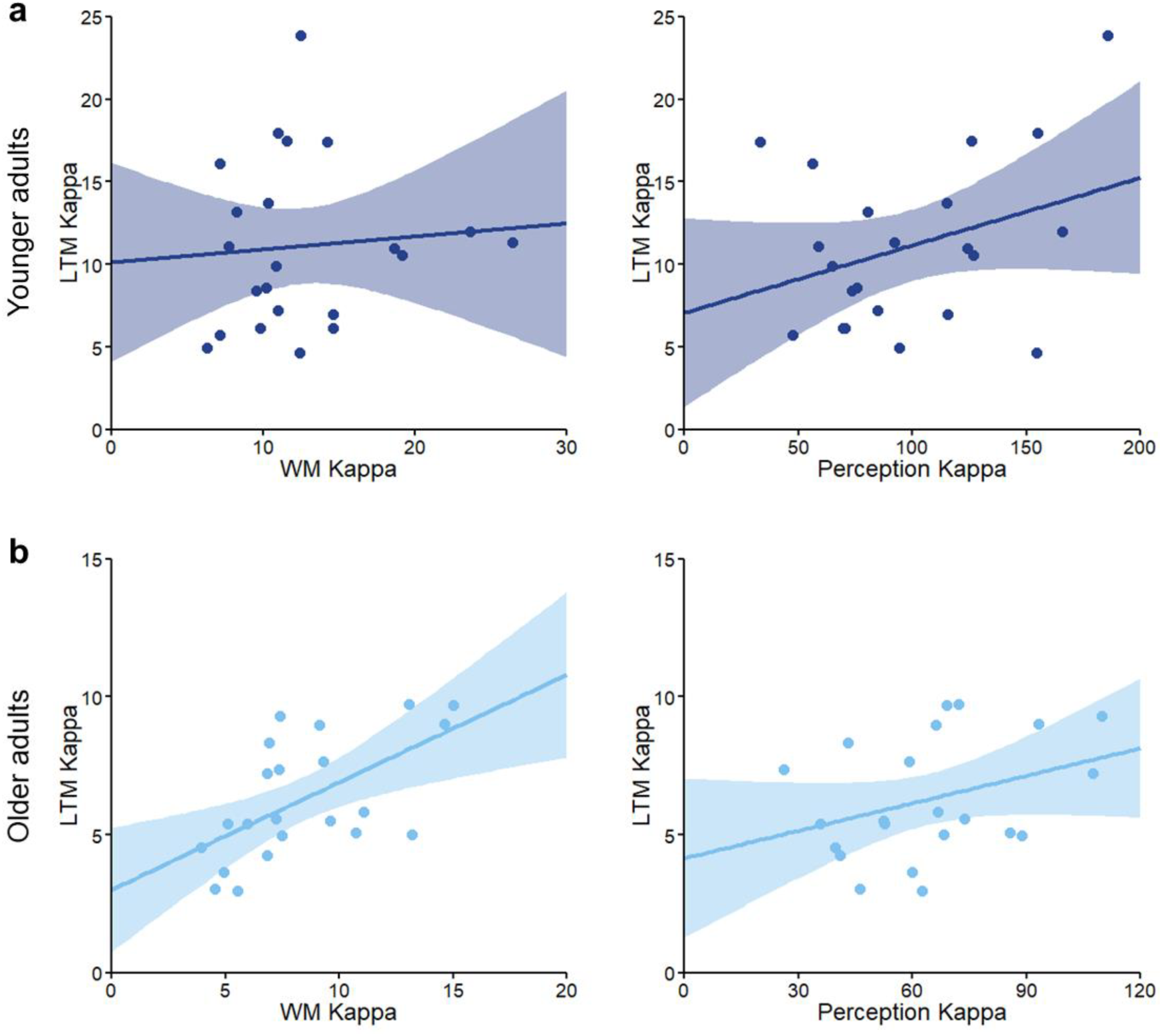
Relationship between LTM and WM precision, and LTM and perceptual precision in the a) younger and b) older groups. Note different scaling of axes between the age groups.

With working memory precision accounting for 35% of variance in the precision of LTM retrieval in the older group, we next examined whether the age-related deficits in LTM precision persisted after controlling for the age-related reductions in the precision of WM retrieval. Critically, after controlling for variability in WM precision, *F*(1, 41) = 10.23, *p* = .003, *partial η^2^* = .20, and after controlling for variability in both WM and perceptual precision, *F*(1, 40) = 6.41, *p* = .015, *partial η^2^* = .14, in an ANCOVA, we still observed significant age-related declines in the precision of LTM retrieval.

## Discussion

In Experiment 3, we examined the degree to which age-related changes in the fidelity of perception and/or WM might contribute to the observed age-related deficits in episodic memory precision. Consistent with findings from Experiments 1 (location) and 2 (location and colour conditions), in the present experiment we observed significant age-related declines in the precision of LTM, but no evidence for age differences in LTM retrieval success. Similarly, the precision of WM and perception were also reduced in the older group, but we did not observe significant age differences in the probability of successfully reporting the correct colour in either of the two tasks. We next assessed the extent to which LTM precision might be constrained by variation in perceptual and WM precision. Despite significant age-related decreases in perceptual precision, perceptual precision did not account for a significant proportion of variance in the precision of LTM in the old or younger adults, and both groups in general performed at a high level on this task. WM precision, on the other hand, was a significant predictor of LTM precision in the older, but not younger adults, suggesting a contribution of decreases in the fidelity of WM to the deficit in LTM precision observed in the older group. However, critically, we still observed age-related declines in the precision of LTM after controlling for variability in the precision of both perception and WM, indicating that the observed age-related reductions in LTM precision could not fully be accounted for by working memory differences.

## General Discussion

The current study sought to better characterise the nature of age-related declines in episodic memory, specifically distinguishing whether age-related changes reflect reduced probability of successfully retrieving information from memory, and/or decreased precision of the retrieved memory representations. Across all three experiments, we consistently observed age-related reductions in the precision of episodic memory retrieval. Declines in mnemonic precision were observed for retrieval of both contextual (Experiments 1 and 2) and item-specific information (Experiments 2 and 3), and persisted after controlling for age-related decreases in the precision of perception and WM (Experiment 3). Reductions in retrieval success, on the other hand, were observed only in the orientation condition in Experiment 2, which was also the condition that resulted in the lowest retrieval success across both age groups (suggesting greater task difficulty). Together, these results highlight reduced precision of memory representations as one factor contributing to age-related episodic memory impairments, and suggest that the success and precision of episodic recollection might be differentially sensitive to age-related cognitive decline.

The current findings of age-related degradation of the precision of episodic memory retrieval are consistent with previous accounts proposing a loss of quality, and specificity, of memory representations in older age (Burke et al., 2018; Goh, 2011; Li, Lindenberger, & Sikström, 2001; Trelle et al., 2017). The observed pattern of reduced precision with preserved retrieval success is in line with studies proposing that while memory for the gist of an event, or stimulus, might be preserved in ageing, the more fine-grained details tend to be lost (Dennis et al., 2007, 2008; Kensinger & Schacter, 1999; Nilakantan et al., 2018). However, whereas previous experiments have predominantly relied on categorical measures of memory success, tasks that emphasise spatial features exclusively, or comparisons between tasks conditions to draw inferences about changes in memory quality, the current study provides a more direct and versatile behavioural measure of memory fidelity, separable from differences in the probability of successful retrieval. The current paradigm has the advantage of allowing the probability of retrieval success and retrieval precision to be estimated from the same data, facilitating the interpretation of results in contrast to inferences drawn from comparisons between different task manipulations. Furthermore, when directly comparing age differences in retrieval success and precision, we observed disproportionate declines in the precision of episodic memory in most task conditions, providing evidence for greater sensitivity of retrieval precision to age-related cognitive decline.

The present continuous report task also permits memory precision of a variety of features to be tested (here, location, orientation, and colour, but in theory, many other features too), enabling the detection of age-related vulnerability across the diverse qualities of mental experiences that characterise episodic remembering. Contrary to the notion of relative sparing of item-memory in older age (Old & Naveh-Benjamin, 2008; Spencer & Raz, 1995), the current findings indicated that the fidelity of both contextual and item-based memory declines in older age. While age-related deficits in item-memory might not be detected in tasks where a relatively unspecific memory of the item can support performance, for example recognition of studied objects as previously encountered (e.g., Chalfonte & Johnson, 1996), tasks requiring retrieval of more specific information of the item, such as discrimination between studied objects and similar lure items, often lead to age-related decreases (e.g., Stark et al., 2013; Trelle et al., 2017), consistent with the current findings of reduced item memory precision in older age. Furthermore, in the current data, age differences in retrieval precision did not significantly vary across the feature conditions, potentially suggesting similar age-related reductions in representational fidelity across different types of information stored in LTM, a hypothesis that future experiments testing precision of other features could confirm.

The findings of age-invariance in the probability of successful memory retrieval in the current experiments contrast with previous reports of age-related declines in episodic recollection success (e.g., Cansino et al., 2018; Simons, Dodson, Bell, & Schacter, 2004). This apparent discrepancy might be explained by age-related decreases on categorical measures of memory success in previous studies being partially attributable to reduced fidelity of the underlying memory representations, rather than a failure to retrieve the representations per se (Nilakantan et al., 2018). For instance, a failure to discriminate between two similar sources of memories, such between two female or two male voices (Simons et al., 2004), could result from a noisier memory representation of the source, leading to selection of the incorrect retrieval response, and thereby reduced retrieval success. In the current experiments, age-related declines in the probability of successful retrieval were observed only in the orientation condition in Experiment 2. Instead of reflecting a feature-specific impairment, it might be that age differences in retrieval success in this condition emerged with increasing task difficulty, consistent with the idea that age differences in cognitive performance are exaggerated when task demands are high (Reuter-Lorenz & Cappell, 2008). The observation that the orientation condition was associated with lower retrieval success than location or colour retrieval across both age groups supports this interpretation. However, future research employing different task difficulty manipulations on the same feature condition is required to evaluate this interpretation.

One concern when investigating age-related memory reductions is that observed differences might be a direct consequence of more general cognitive decline with age, including decreased perceptual abilities (Baltes & Lindenberger, 1997; Lindenberger & Baltes, 1994). The results of Experiment 3 ruled out differences in fine perceptual discrimination or sensorimotor accuracy as a sufficient explanation for the observed deficit in mnemonic precision. Even though older adults were significantly less precise at matching the colour of objects in the perceptual control task, both age groups performed at a high level on this task, and variability in perceptual precision did not significantly account for variability in the precision of long-term memory retrieval in either the young or older adults. Working memory precision, on the other hand, was a significant predictor of long-term memory precision in the older group, indicating that age-related declines in the fidelity of working memory contributed to the deficit observed in long-term memory. The results imply a common factor limiting the fidelity of mnemonic representations in older age both over the short and long term. This is consistent with previous reports highlighting the importance of working memory processes for successful long-term memory encoding (Blumenfeld & Ranganath, 2006; Khader, Jost, Ranganath, & Rösler, 2010), as well as findings implicating working memory as a predictor of episodic memory function in ageing (Bender & Raz, 2012; Hertzog, Dixon, Hultsch, & MacDonald, 2003). However, the effect of working memory on long-term memory observed in the current data was specific to the precision of working memory retrieval, whereas the probability of WM retrieval success did not significantly predict the success, or precision, of long-term memory. Critically, we still observed significant age-related declines in the precision of long-term memory retrieval after controlling for variability in the precision of both perception and working memory. This finding indicates that working memory differences cannot fully account for the observed deficit in episodic memory precision, suggesting additional age-related degradation of information retained in long-term memory.

Previous research has postulated deficient binding of events and features as a key mechanism of age-related decline in episodic memory (Naveh-Benjamin, 2000; Naveh-Benjamin, Hussain, Guez, & Bar-On, 2003), as well as demonstrating age-related increases in binding errors in working memory (e.g., Peich et al., 2013). In the current investigation, we measured binding errors as the probability that participants reported a cued feature of a non-target item from the same study display (e.g., reporting the colour of another object from the same study display as the object tested) (Bays et al., 2009), but found no evidence for binding errors in the older, or younger, group in any of the long term memory tasks or the working memory task (see Supplementary material for model comparison). It might be that in contrast to previous working memory investigations employing simple shape stimuli (e.g., Peich et al., 2013), the object stimuli used in the current experiment resulted in enhanced performance because participants could draw on each object’s rich, semantic representation, on which to bind the individual target features. This hypothesis is in line with previous reports demonstrating a benefit of real-word object stimuli for working memory performance (Brady, Störmer, Alvarez, 2016), as well as preserved ability to utilize semantic information to support memory functioning in older age (Crespo-Garcia, Cantero, & Atienza, 2012; Mohanty, Naveh-Benjamin, & Ratneshwar, 2016; Naveh-Benjamin, Craik, Guez, & Kreuger, 2005). Alternatively, it is also possible that in the present long-term memory tasks, participants made binding errors *across* study displays, potentially driven by semantic or perceptual similarity of the items rather than shared context (i.e., background pictures). Future experiments could distinguish these hypotheses by manipulating the semantic and perceptual relatedness of stimuli both within and across study displays.

The current findings of differential effects of ageing on episodic memory retrieval success and precision imply distinct neurocognitive factors contributing to age-related changes on each component. At the neural level, previous results by Richter, Cooper and colleagues (2016) in younger adults have demonstrated the success and precision of episodic memory retrieval to rely on dissociable brain regions of the core recollection network, with retrieval success associated with activity in the hippocampus, and retrieval precision scaling with activity in the angular gyrus. These findings are consistent with the idea that in response to a retrieval cue, the hippocampus initiates memory retrieval via the process of pattern completion (Norman & O’Reilly, 2003), and might provide a threshold memory signal (Norman, 2010; Yonelinas, 2002), denoting instances in which the cue either succeeds or fails to elicit recollection. Retrieved memories are further reinstated in cortical regions (Bosch, Jehee, Fernandez, & Doeller, 2014; Treves & Rolls, 1994; Wheeler, Petersen, & Buckner, 2000), and the angular gyrus may play a role in online representation of the integrated, episodic, content (Bonnici, Richter, Yazar, & Simons, 2016; Rugg & King, 2018).

Given the putative role of hippocampus and angular gyrus in the success and precision of episodic memory retrieval, respectively, it might be that the behavioural results observed in the present data map onto distinct age-related functional and structural alterations in these two brain regions. Previous investigations have demonstrated diminished episodic recollection effects in the angular gyrus in older adults (e.g., Daselaar, Fleck, Dobbins, Madden, & Cabeza, 2006; Duarte, Graham, & Henson, 2010), potentially contributing to the impoverished precision of retrieved memories observed here. The absence of age differences in the probability of retrieval success in most task conditions in the current study might seem surprising given previous reports of age-related declines in both function (Daselaar et al., 2005; Duverne, Habibi, & Rugg, 2008) and structure (Raz et al., 2005; Walhovd et al., 2011) of the hippocampus. However, not all studies have observed retrieval-related changes in this region in healthy older adults (Persson, Kalpouzos, Nilsson, Ryberg, & Nyberg, 2011; Wang, Johnson, de Chastelaine, Donley, & Rugg, 2016), with some finding a lack of age effects when controlling for reductions in behavioural performance (de Chastelaine, Mattson, Wang, Donley, & Rugg, 2016), highlighting the importance of distinguishing between functional changes due to age and performance.

With the growing ageing population, maintenance of memory abilities in older age is of critical importance from both individual and societal perspectives, emphasising the need for sensitive behavioural markers of early age-related declines. Previous work suggests that tasks requiring reconstruction of studied stimuli may provide a more sensitive measure of age-related memory decline (Clark et al., 2017), highlighting a potential benefit of continuous report measures, such as those used in the current study. In contrast to more traditional, categorical measures of memory performance, fine-grained multi-featural assessment of retrieval with continuous report measures may prove advantageous for early detection of age-related changes in the complex, multifaceted qualitative aspects of memory retrieval. These types of tasks further have the benefit of disentangling the effects of treatments and interventions designed to boost memory functioning in older age on different subcomponents of memory retrieval. Whereas previous work has primarily focused on enhancing the success of memory retrieval, different strategies may be required to ameliorate reductions in retrieval precision, which as indicated by our current data appear to be consistently observed even in healthy older individuals, and are often disproportionate to any changes in retrieval success.

In conclusion, the current study employed continuous retrieval measures to elucidate the mechanisms of age-related changes in episodic memory, identifying age-related declines in the precision of episodic memory representations. Age-related decreases in the fidelity of episodic memory were evident even in the absence of age differences in the probability of successful retrieval, suggesting that this aspect of episodic retrieval might be more sensitive to age-related degradation in the healthy population. Furthermore, age-related declines in mnemonic precision were evident for both item-based and contextual memory retrieval, and were influenced, but not fully explained, by age-related reductions in the precision of working memory. The findings highlight the benefit of continuous report paradigms for revealing the specific basis of memory impairments associated with older age, and call for investigation of the potentially dissociable neural mechanisms underlying age-related changes in the success and precision of episodic memory retrieval.

## Supplementary material

### Model comparison

In each experiment, we compared three alternative models capturing participants’ performance in the continuous report tasks (see Figure S1) (cf., Richter, Cooper et al., 2016). The first model (Model 1), consisted of a von Mises distribution centred at the target feature value alone. This model assumes that on all trials a participant successfully retrieves information about the target feature value, with variable degree of noise. The second model (Model 2), includes both a target von Mises distribution and a circular uniform distribution. In this model, participants’ responses are considered to reflect a combination of variability in target retrieval and random guess responses, where memory retrieval has failed. In addition to the target von Mises and uniform components, the third model (Model 3), includes von Mises distributions centred at the non-target feature values from the same encoding display, incorporating a further component of error: non-target reports, where participants mistakenly report a cued feature value of another item from the same study display (i.e., reporting the location of another object from the same display) (Bays et al., 2009). Consistent with previous results from younger adults (Richter, Cooper et al., 2016), Model 2 fit both younger and older participants’ data better than the two alternative models in each experiment, as indicated by lower Bayesian information criterion (BIC) for this model in comparison to the other two models (see Table S1). Model 2 was therefore fit to participants’ data in all experiments.

**Table S1.**
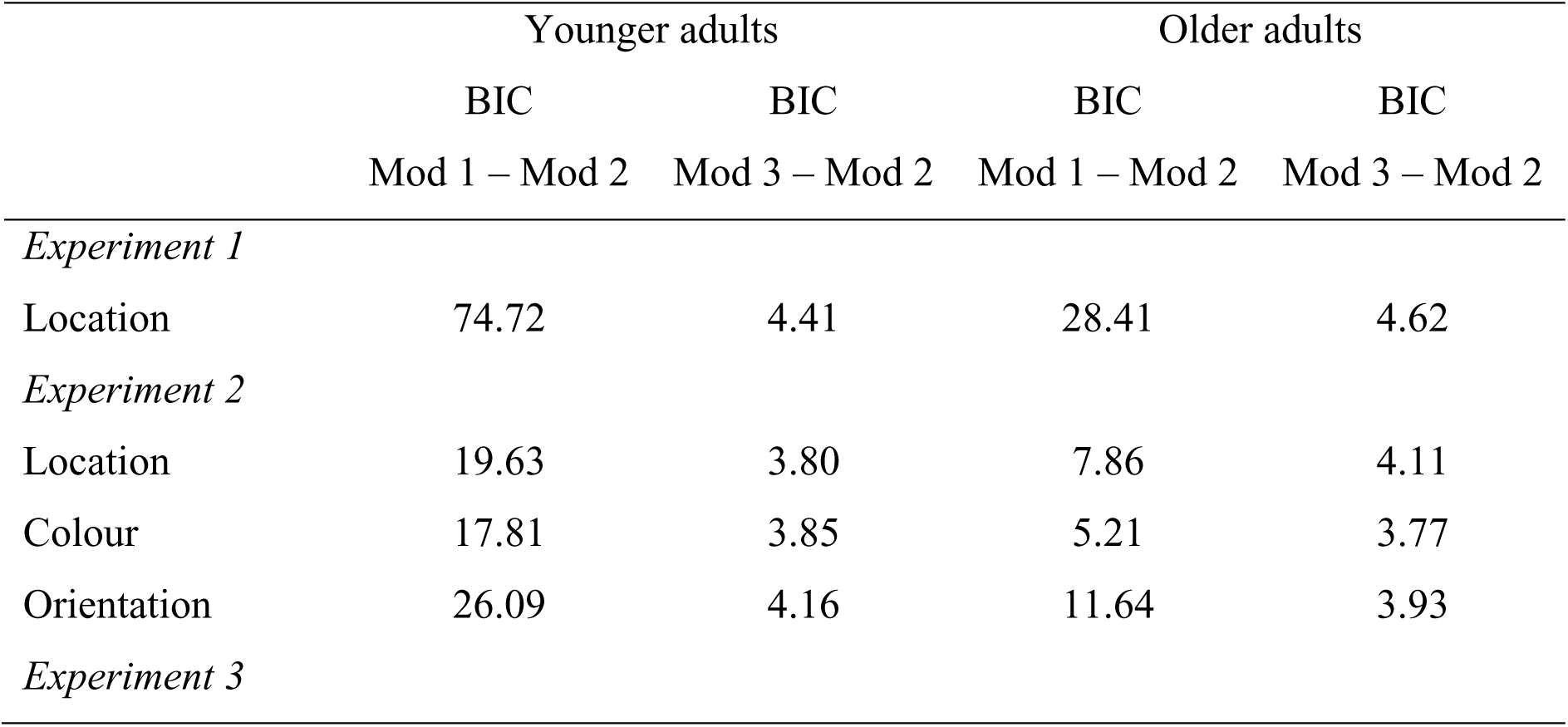
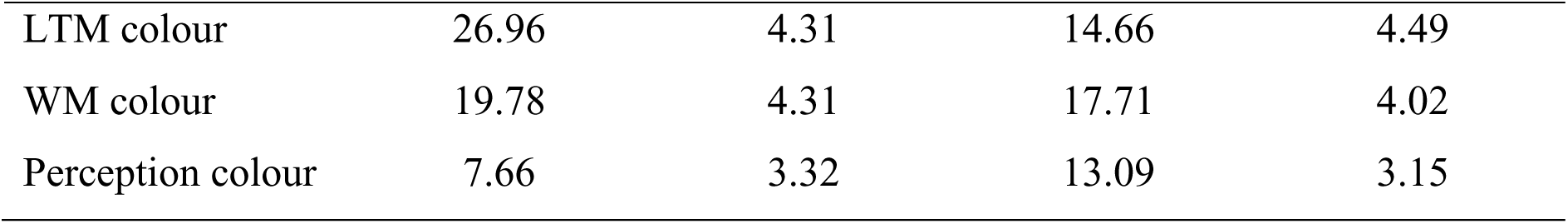
Mean BIC difference between the preferred model (Model 2: uniform and target von Mises) and the two alternative models (Model 1: von Mises only, Model 3: uniform, target von Mises, and non-target von Mises) in each age group.

**Figure S1.**
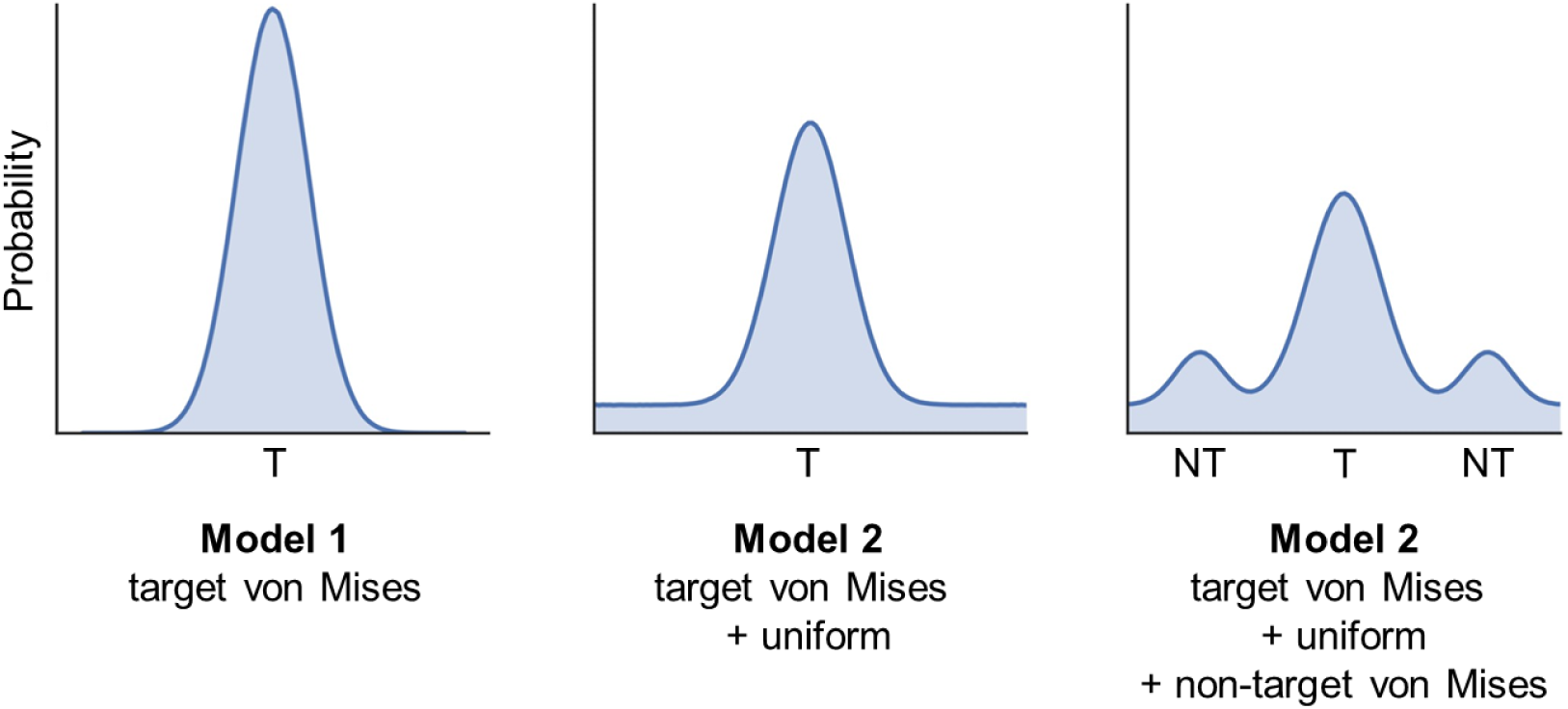
Three alternative models fit to participants’ error data in each experiment. Model 1 includes only a von Mises distribution around the target (T) feature value. Model 2 consists of a target von Mises distribution and a circular uniform distribution. Model 3 includes target von Mises and uniform distributions, as well as von Mises distributions around the non-target (NT) features values from the same encoding display.

### Aggregate analysis

Due to a relatively small number of trials per feature condition in the second experiment, we further replicated our analyses by modelling performance across all trials and participants in each age group and task condition (minimum number of trials per condition via this method was 1920 trials). This approach is less susceptible to noise in parameter estimates in the cases of individual participants with poor performance (cf., Cooper et al., 2017). Maximum likelihood estimates of the probability of retrieval success and the precision of target retrieval in the aggregate analyses are displayed in Table S2. For the aggregate analyses, the statistical significance of the observed age differences on the each parameter estimate were assessed via permutation tests, where participants’ data was remodeled over 1000 iterations of random participant-to-group assignments. To obtain a two-tailed p-value, the absolute observed group difference was compared to the distribution of permuted group differences across the iterations.

The results from aggregate analysis closely mirrored those of the individual participant analysis. In Experiment 1, we observed no significant age differences in the probability of successful memory retrieval (*p* = .407), but a significant decline in retrieval precision (*p* < .001). In Experiment 2, no significant age-related changes in retrieval success were evident in the location (*p* = .890), and colour conditions (*p* = .145), however the older adults performed significantly worse in the orientation condition (*p* = .036). Retrieval precision in Experiment 2, on the other hand, was significantly reduced in the older group in all three feature conditions (location: *p* = .002, colour: *p* < .001, orientation: *p* = .032). In Experiment 3, no significant group differences in the probability of reporting the target colour were observed in the LTM (*p* = .272), WM (*p* = .667), or perception task (*p* = .103), whereas the older adults were significantly less precise in all three tasks (LTM: *p* < .001, WM: *p* < .001, perception: *p* < .001). Findings from the aggregate analyses therefore confirmed the pattern of age-related changes observed when modelling individual participants’ performance.

**Table S2.** Retrieval success (pT) and precision (K) estimated across all trials and participants in each age group and task condition.

|  | Younger adults |  | Older adults |  |
| --- | --- | --- | --- | --- |
| | $pT$ | $K$ | $pT$ | $K$ |
| *Experiment 1* |  |  |  |  |
| Location | 0.82 | 24.30*** | 0.77 | 10.35*** |
| *Experiment 2* |  |  |  |  |
| Location | 0.78 | 11.12** | 0.79 | 6.46** |
| Colour | 0.67 | 8.72*** | 0.59 | 4.26*** |
| Orientation | 0.51* | 12.54* | 0.36* | 8.84* |
| *Experiment 3* |  |  |  |  |
| LTM colour | 0.82 | 10.00*** | 0.77 | 5.68*** |
| WM colour | 0.91 | 11.73*** | 0.90 | 7.29*** |
| Perception colour | 0.99 | 88.45*** | 0.98 | 52.60*** |

*Note.* Significant between group differences are indicated by asterisks next to each groups parameter estimates (\* $p < .050$ , \*\* $p < .010$ , \*\*\* $p < .001$ ).

## Acknowledgements

This study was funded by BBSRC grant BB/L02263X/1 and James S. McDonnell Foundation Scholar Award #220020333, and was carried out within the University of Cambridge Behavioural and Clinical Neuroscience Institute, funded by a joint award from the Medical Research Council and the Wellcome Trust. We are grateful to Paul Bays for valuable advice, and to Lowri Foster Davies and Stephanie Ngai for assistance with data collection.

